# Harmonizing terrestrial carbon cycle observations over CONUS NEON sites: Assessing the information contributions of multiple data constraints

**DOI:** 10.1101/2025.06.11.659109

**Authors:** Dongchen Zhang, Qianyu Li, Alexis Helgeson, Shawn P. Serbin, Michael Dietze

## Abstract

Accurate inventories of terrestrial carbon pools and fluxes are crucial for understanding ecosystem processes, tracking climate change impacts, and meeting the monitoring, reporting, and verification (MRV) requirements in international treaties and voluntary carbon markets. In meeting this need, the fusion of process-based modeling, field data, and remote sensing observations has the potential to provide more accurate and precise estimates than each alone. However, as the number of data constraints on a system increases, different sources of information can interact with each other in complex ways across space, time, and processes. In this study, we undertake a value-of-information analysis to assess the contribution of different observations to reducing carbon cycle uncertainties across pools, fluxes, and spatial domains within the PEcAn carbon cycle data assimilation system. We used a novel block-based Tobit Gamma Ensemble Filter to assimilate four synergistic data constraints, MODIS leaf area index, Landtrendr aboveground biomass, SMAP soil moisture, and SoilGrids soil organic C, into a process-based ecosystem model (SIPNET) at 39 National Ecological Observatory Network sites across the contiguous U.S. from 2012 to 2021. Results showed that not only did we greatly reduce uncertainty among the directly constrained pools but many observations were able to share information across variables and space. These indirect constraints helped identify synergies and conflicts among data streams and across space, which provides insights for further constraining carbon inventories. Overall, soil carbon remains the largest source of uncertainty in the overall carbon budget due to both its large size and limited observational constraints.

## 1. Introduction

The accurate monitoring, reporting, and verification (MRV) of terrestrial carbon (C) pools and fluxes have gained considerable attention as the land biosphere is a significant sink of carbon from the atmosphere that helps mitigate the effects of global climate change (Le Quéré et al., 2015). However, constraining estimates of C pools and fluxes are challenging as terrestrial ecosystems are highly variable across both space and time and can often become a net source of C due to human activity, urbanization, natural disturbances, and extreme climate events (Battle et al., 2000). From an ecological perspective, the uncertainty in the terrestrial C cycle also comes from complex interactions among organisms and between organisms and nutrients, water, and climate, which makes land C cycling challenging to measure and quantify. This combination of high variability and system complexity means that the terrestrial C cycle is one of the least constrained components of the global carbon budget (Ballantyne et al., 2017).

Numerous efforts have been made to reduce MRV uncertainties in the terrestrial C cycle using various combinations of ground-based measurements, remote sensing, and modeling (Hurtt, et al., 2022). In many ways, ground-based approaches remain the “gold standard” but these approaches are labor-intensive, hard to scale, and are rarely able to measure the full suite of all relevant pools and fluxes (Tang et al., 2018; Pan et al., 2011; Xiao et al., 2014). Remote sensing observations are becoming increasingly important in C monitoring studies as a way to scale up ground-based measurements based on a variety of sensing technologies covering different spatial and temporal domains and resolutions (Hurtt et al., 2022). However, these remotely-sensed observations are used to produce individual data products that estimate a single pool or flux, and thus each only tells a partial story about the C cycle (Asner & Ollinger, 2009; Wang et al., 2021; O’Sullivan et al., 2022). While many such products exist, it can be difficult to reconcile different data products into a harmonized understanding of the C cycle as a whole in a way that correctly accounts for their uncertainties and the pools and fluxes that cannot be observed directly (Kennedy et al., 2024; Wikle, 2015).

Terrestrial biosphere models predict C pools and fluxes based on mathematical approximations of key processes, including photosynthesis (Farquhar et al., 1980; Ball et al., 1987), carbon allocation (Litton et al., 2007), respiration (Richardson et al., 2005), turnover, and decomposition (Berg et al., 2000). Not only do these models embed our understanding and hypotheses about how the C cycle functions, but they also provide partitioned and balanced ecosystem C budgets across consistent spatial and temporal scales. Recent trends in land carbon modeling have been toward models increasingly calibrated and/or benchmarked against remotely sensed observations and the integrated networks such as ILAMB (International Land Mode Benchmarking, Hoffman et al., 2017), because of the gridded data layers they provided at continuous and large spatial and temporal scales (Fer et al., 2021). However, even calibrated and validated terrestrial biosphere models are simulations of the ecosystem, not a record of what actually happened at a specific time and place, and are limited by our imperfect understanding of complex ecological processes (Xu et al., 2020) and the presence of stochasticity in real-world events (e.g., disturbances). Overall, there is an urgent need to better integrate the information provided by ground observations, remote sensing, and models to improve both C monitoring and our understanding of ecological processes and functions.

State Data assimilation (SDA) provides a potential path forward for addressing the challenges facing C accounting (Baatz et al., 2021). SDA is a statistical technique that integrates observations into models to constrain a model’s state variables (e.g., pools and fluxes) rather than its parameters (a.k.a. calibration). It assumes that neither models nor observations are sufficient to represent the dynamics of a system and that combining observations and models can be more accurate and precise than either alone (Williams et al., 2005). In its simplest form, SDA approaches improving model predictions by “nudging” them towards reality. But in doing so SDA shares information across multiple data streams, space, and ecological processes such that unobserved locations or ecological variables can be inferred from data-rich regions and easily observed variables (Dietze, 2017). As such, the model-based fusion of data streams can integrate sparse or discontinuous data, and can be used to provide coverage across space and time to make better estimates and inferences than any single data stream can provide alone. When run post-hoc (as opposed to part of an automated forecast), this model-data fusion product is commonly referred to as a “reanalysis”. Given the strengths of SDA, these approaches have been widely applied in the atmospheric sciences for decades (Daley, 1993) and have more recently been adapted to hydrological and ecological studies (Dong et al., 2016).

SDA has increasingly been employed to constrain estimates of the land C cycle. Leaf area index (LAI) data has been used extensively in land C SDA studies (Mao et al., 2017; Li et al., 2019; Ramos et al., 2018; Chen & Tao., 2020; Kumar et al., 2019; Revill et al., 2013) due to its importance in modeling (e.g., gross primary productivity (GPP), evapotranspiration (ET)) and its wide availability from both remote sensing and ground-based approaches (MODIS LAI; tower-based LAI). In addition to assimilating LAI observations, an increasing number of land C SDA studies have begun to assimilate additional data streams. For example, assimilating both LAI and soil moisture tends to improve predictions of vegetation dynamics and their interactions with the water cycle (Fairbairn et al., 2017; Sawada et al., 2015; Sabater et al., 2008; Albergel et al., 2010). Other studies have assimilated: 1) LAI and AGB, leading to more accurate predictions of soil organic carbon (SOC) (Dokoohaki et al., 2022); 2) LAI and fraction of absorbed photosynthetically active radiation (faPAR) to improve phenological predictions (Viskari et al., 2015); and 3) soil moisture and CO2 measurements with improved estimations of land C fluxes (Wu et al., 2020).

These SDA analyses have proven effective in reducing both the residual errors between the carbon cycling reanalysis and the correspondent observations and the predictive uncertainty of the reanalysis itself. However, to date, SDA has only been applied to a subset of ecological processes within terrestrial biosphere models using a limited number of data types. We are unaware of any land C SDA studies that have assimilated more than two data types for the purpose of constraining C stocks instead of model parameters. Because of this, SDA approaches have yet to be extended to harmonize observations of all the major ecosystem C pools (e.g., leaf, stem/root, soil) simultaneously. This raises concerns that, like with many observational approaches, major C pools may be inferred as the “remainder” of the overall C budget. Therefore, there is a need to assimilate data on all major C pools to close large-scale C budgets.

To begin to address this need, we deployed a SDA framework at a smaller spatial scale – across the 39 contiguous United States (CONUS) National Ecological Observatory Network (NEON) sites – using four data types: MODIS LAI, Landtrendr AGB, SMAP Soil moisture, and SoilGrids SOC. As a proof-of-concept, this system was run from 2012 to 2021 with an annual assimilation of new data and a sub-daily model timestep. The NEON sites provide a broad, statistically rigorous sampling of ecosystem types, while the assimilation itself focuses on data sources available across a wide spatial extent.

From the SDA perspective, multiple pieces of information can be combined by borrowing strength both across variables (e.g., root biomass is correlated with stem biomass, GPP is correlated with LAI) and across space (e.g., knowing soil C at all the sites within 100 km could help better predict soil C at a focal location). While using SDA provides an important opportunity to reconcile information about key C and water pools, it also brings new science questions and challenges regarding how information is being shared across the system: 1) How well can carbon and water observations be reconciled with terrestrial biosphere models? What are the major impacts of data assimilation on overall carbon and water budgets and their uncertainty? 2) What data sources have the greatest impact on constraining the C cycle directly (i.e., when modeled and measured C pools are equivalent) for each C pool? 3) Which indirect constraints (i.e., based on model covariances) are strongest across pools and between pools and fluxes? 4) Which indirect constraints are strongest across space, and how quickly does information sharing drop off with distance? 5) Are there places where different information sources conflict?

To address these questions, we performed a value-of-information (VOI) analysis, which aims to quantify the potential benefits of assimilating additional data to improve the understanding of the uncertainties for the entire system across observations and space. Specifically, we assessed the relative change in assimilation uncertainty for all major C and water pools/fluxes when different data constraints were added or removed. In addition, a spatial VOI was employed to assess the relative uncertainty reduction as we increased the spatial distances considered when calculating across-site covariances. By knowing the contributions of different observations to the uncertainty of SDA outcomes, VOI analysis can help identify key data sources that contribute the most information to understanding C budgets and ecosystem functioning. This information also helps us identify current data gaps regarding when, where, and which data types are most beneficial for reducing C MRV uncertainty, and thus to prioritizing future data collection.

## 2. Methods

### 2.1. Study Area and Data

NEON provides continental-scale long-term measurements through a combination of intensive fieldwork, automated sensors, and airborne remote sensing observations with sites selected to represent the ecological characteristics within twenty ecoclimatic domains across the United States (Figure 1). The well-stratified NEON site selection facilitates the analysis of spatial-temporal patterns of ecological processes. In this study, we used the 39 terrestrial NEON sites that cover the contiguous U.S. (CONUS) region. We classified the 39 CONUS NEON sites into three plant functional types (PFTs) based on the National Land Cover Database (NLCD) (Homer et al., 2012): coniferous forest, deciduous forest, and non-woody systems (grasslands, croplands, etc.).

**Figure 1.**
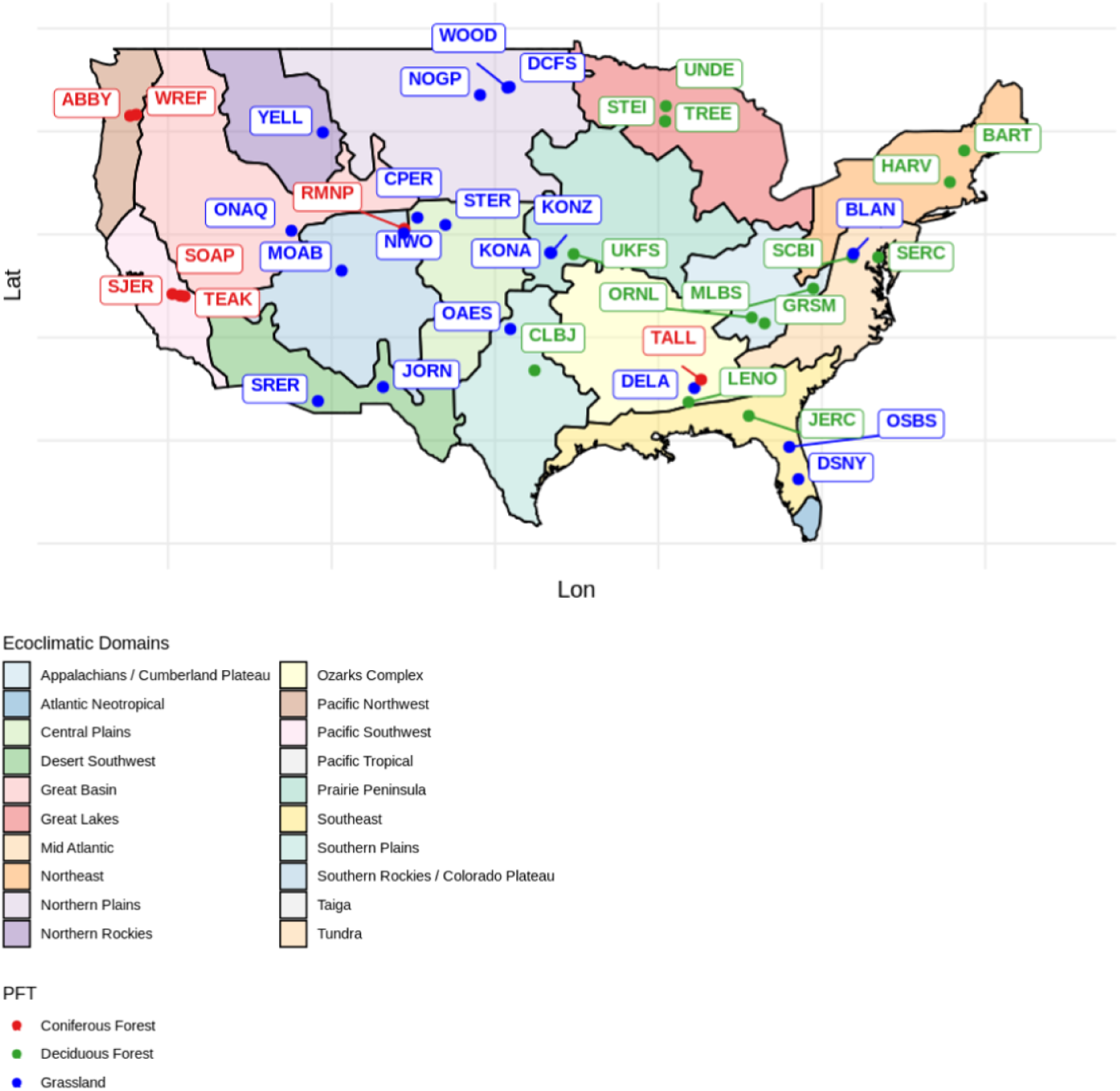
Map of CONUS NEON Sites across Ecoclimatic Domains (level 1 EPA ecoregion map). Sites are labeled with their site codes and colored by their plant functional types.

We selected the temporal domain from 2012 to 2021, given that initial NEON prototype measurements started around 2012 and reached an operational state in 2019. We used three types of datasets in this study (Table 1): 1) Meteorological drivers; 2) carbon pool initial conditions (see supplement 1 for more details); 3) remote sensing observations and other gridded data products that are being assimilated. SIPNET simulations were driven with ERA5 reanalysis, a 30km product with a 3-hour temporal resolution, and ten ensemble members to capture driver uncertainty (Hersbach et al., 2023). Model vegetation and soil C pools were initialized based on NEON site-level observations, which were resampled to capture initial condition uncertainty through 100 ensemble members (see supplement 1). Assimilated date constraints were subset to the years being assimilated, to the extent that each data product is available (e.g., SMAP is only available after 2015, Landtrendr AGB stops at 2017), and QA/QC filtered based on provided metadata (e.g., QA/QC band in MODIS LAI products). For products with sub-annual resolutions (8-day MODIS LAI, 3-day SMAP soil moisture; see Table 1) data were further subset to a window (60 days for LAI and SMAP) around the annual assimilation date (July 15th), selecting the closest valid observation to the assimilation date.

**Table 1.** Datasets used in this study. Here, we summarize the input datasets within the SDA framework, their spatial and temporal resolutions, and abbreviations used throughout the manuscript (Note that the NEON initial conditions differ in their dates, which depends on the earliest available measurements for each data).

| Data Type | Data Name | Period | Source | Spatial Resolution | Temporal Resolution |
| --- | --- | --- | --- | --- | --- |
| Meteorological Driver | ERA5 | 2012 - 2021 | 10.24381/cds.adbb2d47 | 30km | 3-Hour |
| Initial Condition | Leaf Biomass | >= 2015 | 10.48443/kbnv-bz87 | 20 - 40m | Yearly |
| Initial Condition | Wood Biomass | >= 2015 | 10.48443/73zn-k414 | 20 - 40m | Yearly |
| Initial Condition | Soil Organic Carbon (SOC) | >= 2015 | 10.48443/t70z-np08;<br>10.48443/0phb-j505 | 20 - 40m | Yearly |
| Observation | LAI | 2012 - 2021 | 10.5067/MODIS/MOD15A3H.006 | 500m | 8-day |
| Observation | Aboveground Biomass (AGB) | 2012 - 2017 | 10.1088/1748-9326/aa9d9e | 30m | 1-year |
| Observation | Soil Moisture Profile (SMP) | 2015 - 2021 | 10.5067/LWJ6TF5SZRG3 | 9skm | 3-day |
| Observation | Soil Organic Carbon (SOC) | 2012 - 2021 | 10.5194/soil-7-217-2021 | 250m | NA |

### 2.2. Ecosystem Model

We used the Simplified Photosynthesis and Evapotranspiration Model (SIPNET) (version r136, https://github.com/PecanProject/sipnet.git), a simple pool-based process model, as the scaffold for making forward predictions of all C pools and fluxes and reconciling predictions with observations (Braswell et al., 2005). SIPNET was selected based on its low computational cost and its simplified process representations of coupled carbon and water cycling, which facilitates interpretability.

SIPNET contains four vegetation carbon pools (leaf, stem, coarse roots, fine roots) and one soil carbon pool (Figure 2). Photosynthesis is calculated using a light-use-efficiency approach that is a function of maximum photosynthesis, LAI, and photosynthetically active radiation, with modifiers for air temperature, vapor pressure deficit, and soil moisture. SIPNET also has a simple representation of vegetation C allocation based on fixed fractions to each plant carbon pool. For deciduous systems, leaf phenology is controlled by a simple growing degree day function. Losses from plant carbon pools are based on fixed turnover rates and maintenance respiration, which is a Q10 function of temperature. Soil carbon is increased by wood and leaf litter and reduced by soil respiration. Finally, soil moisture is modeled using a simple bucket-model approach with precipitation as input and evaporation, transportation, and drainage as outputs.

**Figure 2.**
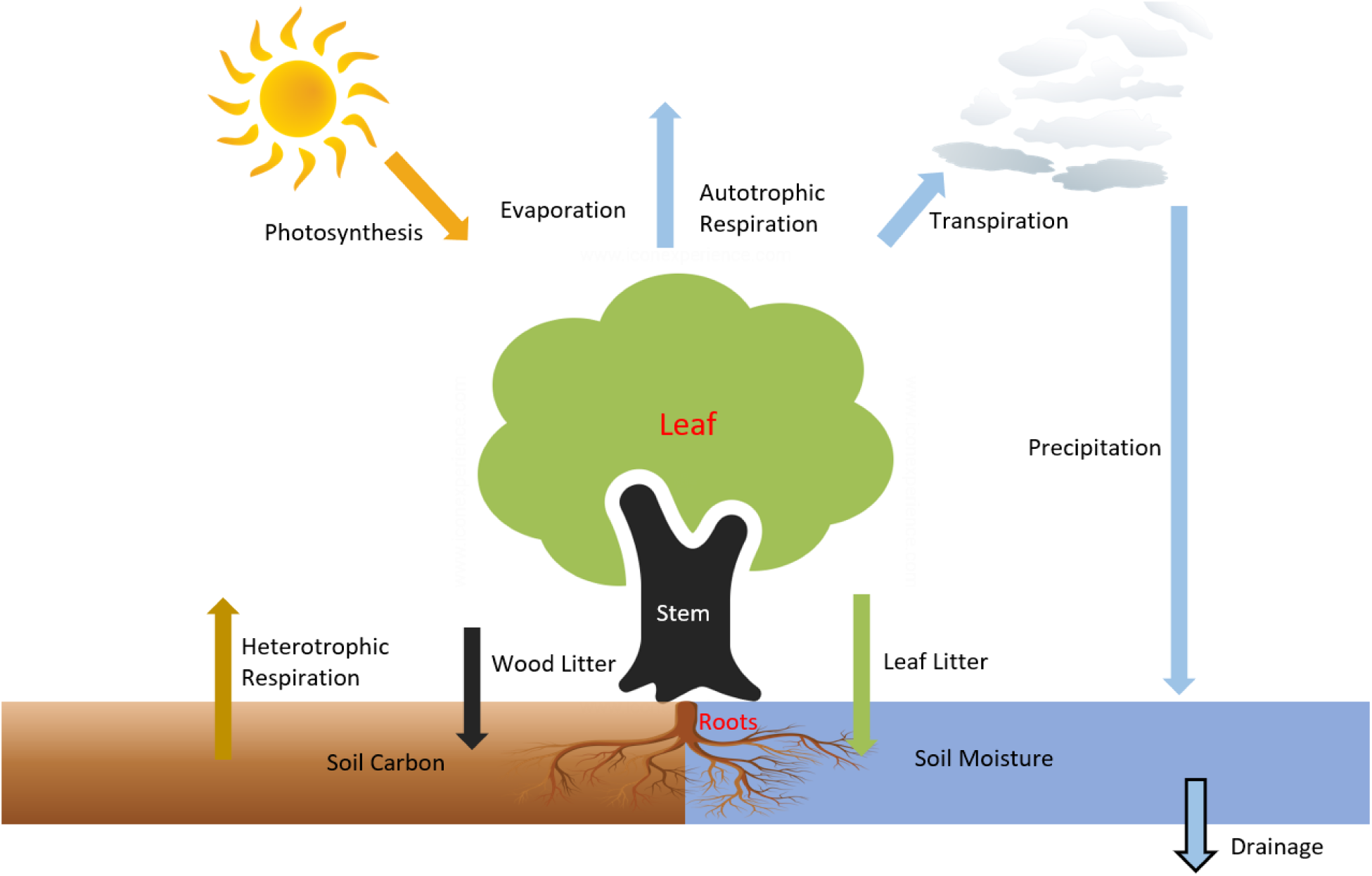
Flowchart of the SIPNET model. The model includes four vegetation carbon pools (leaf, stem, coarse roots, fine roots) and one soil carbon pool. Photosynthesis adds carbon into the system, which is then allocated to the four vegetation pools, while autotrophic and heterotrophic respiration remove carbon from the vegetation and soil carbon pools, respectively. Vegetation pool turnover (a.k.a. litter) adds to soil carbon, and precipitation adds soil moisture, the latter of which is removed by transpiration, evaporation, and drainage.

Model parameter ensembles were sampled from PFT-level joint posterior parameter distributions generated through previous calibrations conducted using a Hierarchical Parameter Data Assimilation (HPDA) approach against the carbon and water flux measurements from 22 Ameriflux towers (Dokoohaki et al., 2022; Fer et al.,2018, 2021).

### 2.4. Model-Data Fusion - The State Data Assimilation (SDA) Approach

#### 2.4.1. Tobit Gamma Ensemble Filter (TGEnF)

In this study, we used a state data assimilation (SDA) approach called the Tobit Gamma Ensemble Filter (TGEnF), a variation of the Tobit Wishart Ensemble Filter (TWEnF) (Dokoohaki et al., 2022), to statistically harmonize the observations and the model forecasts. Similar to other ensemble-based assimilation approaches (e.g., Ensemble Kalman Filter (Evensen, 2003)), this approach begins by making an ensemble prediction of the modeled pools and fluxes, adds in model process error, and then uses this ensemble mean and covariance to parameterize a multivariate Bayesian prior probability distribution. Bayes’s theorem is then used to reconcile this prior forecast with a multivariate distribution of observation and their observation errors (a.k.a. a statistical Likelihood function) to generate an updated posterior distribution that harmonizes the model and data. Unlike other ensemble approaches, TGEnF and TWEnF include an iterative statistical updating of the process error between the model and the unobserved latent state variables. The TGEnF method differs from the TWEnF in that it uses a Normal-Gamma conjugacy between the priors of the process error and the Likelihood instead of the Wishart-Multivariate Normal conjugacy in the TWEnF. Because of this TGEnF doesn’t account for the covariance among model process errors, but the computational demand is considerably lower (because process errors are estimated independently) and the convergence of the Bayesian Markov chain Monte Carlo (MCMC) sampling degrades less as the dimensionality of the assimilation (39 sites × 4 state variables) increases, which makes our multisite-multivariable analysis possible. Additional details on the TGEnF, including information about spatial localization (i.e., the range over which spatial covariances are calculated) and block-based SDA, can be found in supplement 2.

Our overall SDA framework is shown in Figure 4. We first ran 100 SIPNET model ensemble simulations per site, each one year in duration, by sampling inputs from the NEON-based initial conditions (ICs), the ERA5 meteorological drivers, and the PFT-level calibrated posterior parameter distributions. Then, we assimilated the observations into the ensemble forecasts using the TGEnF, the outputs of which become the ICs for the next round of one year SIPNET forecasts. This workflow is implemented within the Predictive Ecosystem Analyzer (PEcAn) model-data informatics system (Dokoohaki et al., 2022; Fer et al., 2018; LeBauer et al., 2013; Raiho et al., 2020), an open-source community cyberinfrastructure project (pecanproject.org, https://github.com/pecanProject/pecan/). Within PEcAn, the TGEnF is implemented using NIMBLE v0.12.1 (de Valpine et al., 2017).

**Figure 4.**
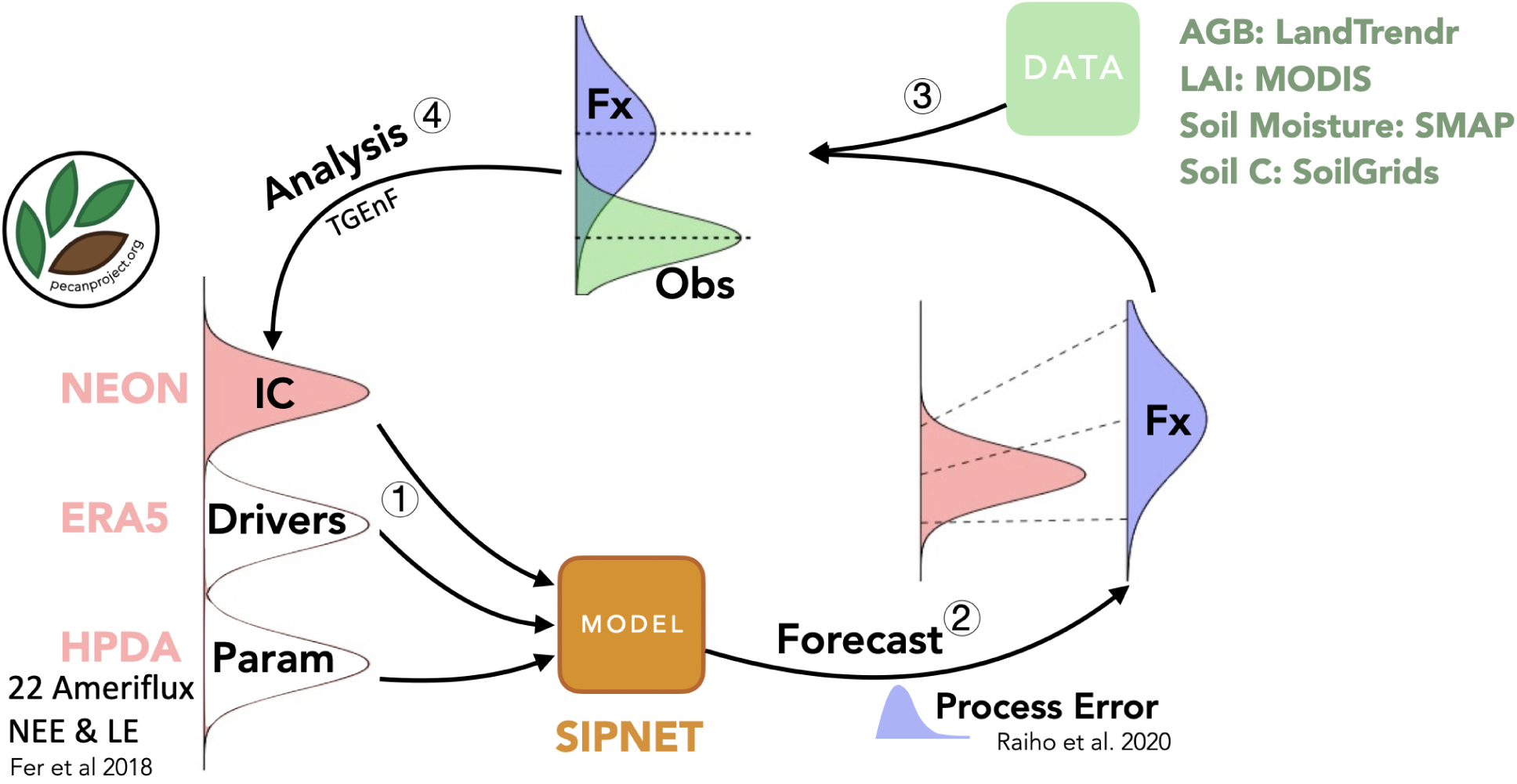
Workflow of the SDA framework. (1) Sample ensemble members from the distribution of data inputs, including NEON initial conditions, ERA5 meteorology drivers, and calibrated parameters. (2) Run an ensemble of the SIPNET model to produce forecasts of key ecological states. (3) As new data become available for each timestep, (4) assimilate the four data streams (AGB, LAI, soil moisture, and soil carbon) using the TGEnF algorithm, producing harmonized estimates of the carbon and water pools and fluxes that can be used as ICs for the next model forecast.

### 2.5. Experiments and Analysis

#### 2.5.1. Definitions of SDA Experiments

Prior to running the SDA, we assessed our initial conditions by comparing the ICs of three carbon pools to the remotely sensed observations (see supplement 3). Because Landtrendr AGB is available at a higher spatial resolution (30m) than both the field data and the other data sources being assimilated, we also analyzed the effect of varying the Landtrendr search radius (from 30m to 150m) on the correlation between the Landtrendr aboveground biomass and the ICs.

The SDA algorithm was then run simultaneously across all 39 NEON sites using all available data constraints (“full run”), producing harmonized spatiotemporal reanalysis estimates of all carbon pools and fluxes, and their uncertainties, including covariances across variables and across space (Baatz et al., 2021). This was then compared to a “free run” of the SIPNET model at each site without any SDA data constraints. In detail, for each run type *r* (in this case *r* ∈ *{free run, full run}*), we produce reanalysis results consisting of *N* = 100 ensemble members for the variable *v* (*v* ∈ *{leaf, wood, soil, root Carbon}*), at site *s*, and time *t*,

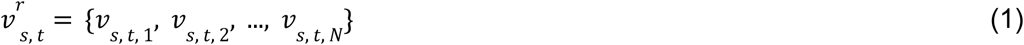

To summarize the spatial variation in carbon pools and associated uncertainties for the free run and full run, we calculate the mean and variance across the whole ensemble space for the run type *r* and variable *v,* at the site *s,* and time *t* according to their sample-based estimators:

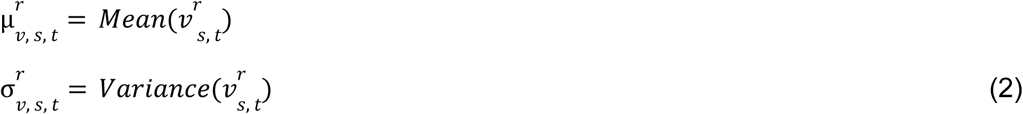

Likewise, the ensemble-based nature of this data product also allows us to easily calculate covariances across variables, space, and time based on sample statistics. Thus, the more generalized standard deviation (SD) and variance calculations across time *T* and space *S* can be written as:

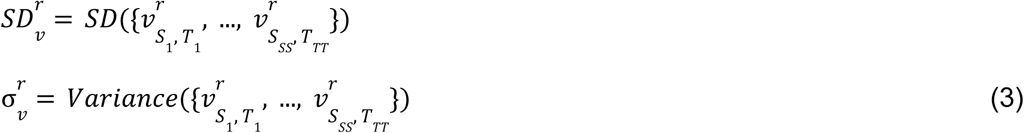

Where *SS* and *TT* represent the length of selected sites *S* and times *T*.

#### 2.5.2. Direct and Indirect VOI Analysis

To assess the value of information (VOI) associated with different data sources and spatial scales, we performed a series of data assimilation experiments with different combinations of observations. One set of experiments involved adding each data constraint individually to the unconstrained “free” run, while the other set involved removing each data constraint individually from the “full” assimilation run (all four constraints). The details of these additional experiments (run types) are listed in Table 2. Based on equation (2), the calculations of VOI for the variable *v* are based on the relative variance increment (in %) between each experiment *r*_2_ (where we added/removed observations of MODIS, Landtrendr, SMAP, or SoilGrids to/from the free run/full run) and the free run or four-observation full run *r*_1_:

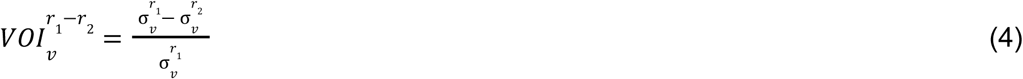

**Table 2.** Lists of SDA experiments summarized by their names, purposes, data streams, and spatial scales.

| Experiment name | Analysis purpose | Data Assimilated | Localization Scale (Km) |
| --- | --- | --- | --- |
| Full Run | Across-variable & Across-space VOI | AGB, LAI, SMP, SOC | 0 |
| -AGB | Across-variable VOI | LAI, SMP, SOC | 0 |
| -LAI | Across-variable VOI | AGB, SMP, SOC | 0 |
| -SMP | Across-variable VOI | AGB, LAI, SOC | 0 |
| -SOC | Across-variable VOI | AGB, LAI, SMP | 0 |
| Free Run | Across-variable VOI | none | 0 |
| +AGB | Across-variable VOI | AGB | 0 |
| +LAI | Across-variable VOI | LAI | 0 |
| +SMP | Across-variable VOI | SMP | 0 |
| +SOC | Across-variable VOI | SOC | 0 |
| 100km Run | Across-space VOI | AGB, LAI, SMP, SOC | 100 |
| 200km Run | Across-space VOI | AGB, LAI, SMP, SOC | 200 |
| 300km Run | Across-space VOI | AGB, LAI, SMP, SOC | 300 |
| 400km Run | Across-space VOI | AGB, LAI, SMP, SOC | 400 |
| 500km Run | Across-space VOI | AGB, LAI, SMP, SOC | 500 |

Similarly, the VOIs for fluxes were calculated based on the predictive variances in the cumulative annual fluxes.

To better understand the correlations that drive specific patterns in both direct and indirect constraints, we also performed a cross-variable VOI analysis to understand the VOI contributed from the variable *v*_1_ to the changes in uncertainty of the variable *v*_2_ across all sites and times between each experiment *r*_2_ and the free run or four-observation full run *r*_1_.

To estimate the VOI for the overall C budget, we calculated the total variance of the variable *v* for the run type *r* across all NEON sites and time points based on Equation (3).

While SDA generally reduces overall uncertainties, it is also possible for VOIs to identify instances when the SDA can’t simultaneously reconcile two data streams with the model. To understand the mechanisms that drive those patterns, we developed a relative change analysis, which compares the direct changes in the mean of a specific variable within the SDA, 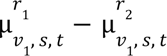 (e.g., the changes of LAI associated with adding or subtracting LAI observations), to the changes in the uncertainties of a second variable, 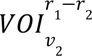 (e.g., GPP).

#### 2.5.3. Residual Error Analysis

In addition to analyzing the posterior uncertainties in the data assimilation, this study also evaluated the residual errors between the assimilation and the held out observations across different SDA experiments (Table 2). Because in the VOI analysis each data stream was sequentially held out of the full assimilation, the residual errors can be interpreted as a cross-validation error. Specifically, the mean absolute error (MAE) of the run type *r* and variable *v* at site *s*, and time *t* across *N* ensemble members for analysis means 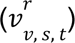 can be written as:

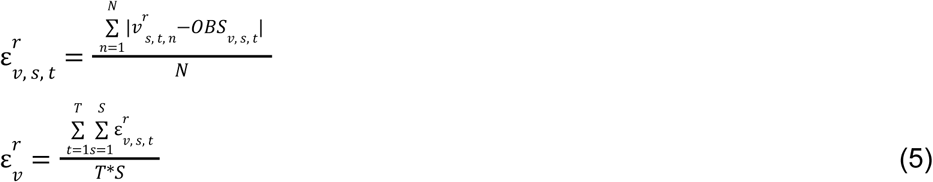

Where *OBS_*v*, *s*, *t*_* is the observation of the variable *v*, at site *s*, and time *t* and 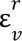 is the average MAE across time and sites.

#### 2.5.4. Spatial VOI Analysis

Because a key feature of SDA is the ability to borrow strength across space, another central aspect that needs to be evaluated is how much uncertainty is reduced as we increase the spatial range of the localization (i.e., distance over which spatial covariances are calculated) (See supplement 2). To do so we generated a series of experiments (See Table 2), relative to the full run, where spatial localization varied between 0 km (which is the same as the full run) to 500 km.Spatial VOI analysis was then conducted to assess the rate at which data constraints decayed as a function of distance. To understand the spatial patterns of uncertainty reduction for different PFTs and variables, the spatial VOI was further visualized by mapping the slope of site-level linear regressions between the variance of the variable *v* and spatial distances *d*, where negative slopes indicate a decrease in uncertainty as the localization scale is increased (i.e., additional borrowing of strength spatially).

## 3. Results

### 3.1. Carbon budgets for CONUS NEON Sites

To systematically describe the carbon density at the site scale before and after assimilating observations, we visualize four carbon pools (leaf, wood, root, and soil carbon) in 2021 between the free run and 500-km run (full run with four observations and 500km localization) (Figure 8). After assimilating observations into the system, the averaged C density across all NEON sites drops from 98.7 kg/m2 to 51.5 kg/m2. For different C budgets, the site-level budgets tended to have less soil organic carbon (from 32.1 to 18.9 kg/m2) overall, (from 36.3 to 20.0 kg/m2) for grass sites, (31.8 to 18.9 kg/m2) for deciduous forest sites, and (22.0 to 15.8 kg/m2) for coniferous evergreen forest sites. The forest sites also tended to have more aboveground biomass (5.4 to 8.0 kg/m2) overall, (7.2 to 14.2 kg/m2) for coniferous evergreen forest sites, and (7.1 to 10.5 kg/m2) for deciduous forest sites.

**Figure 8.**
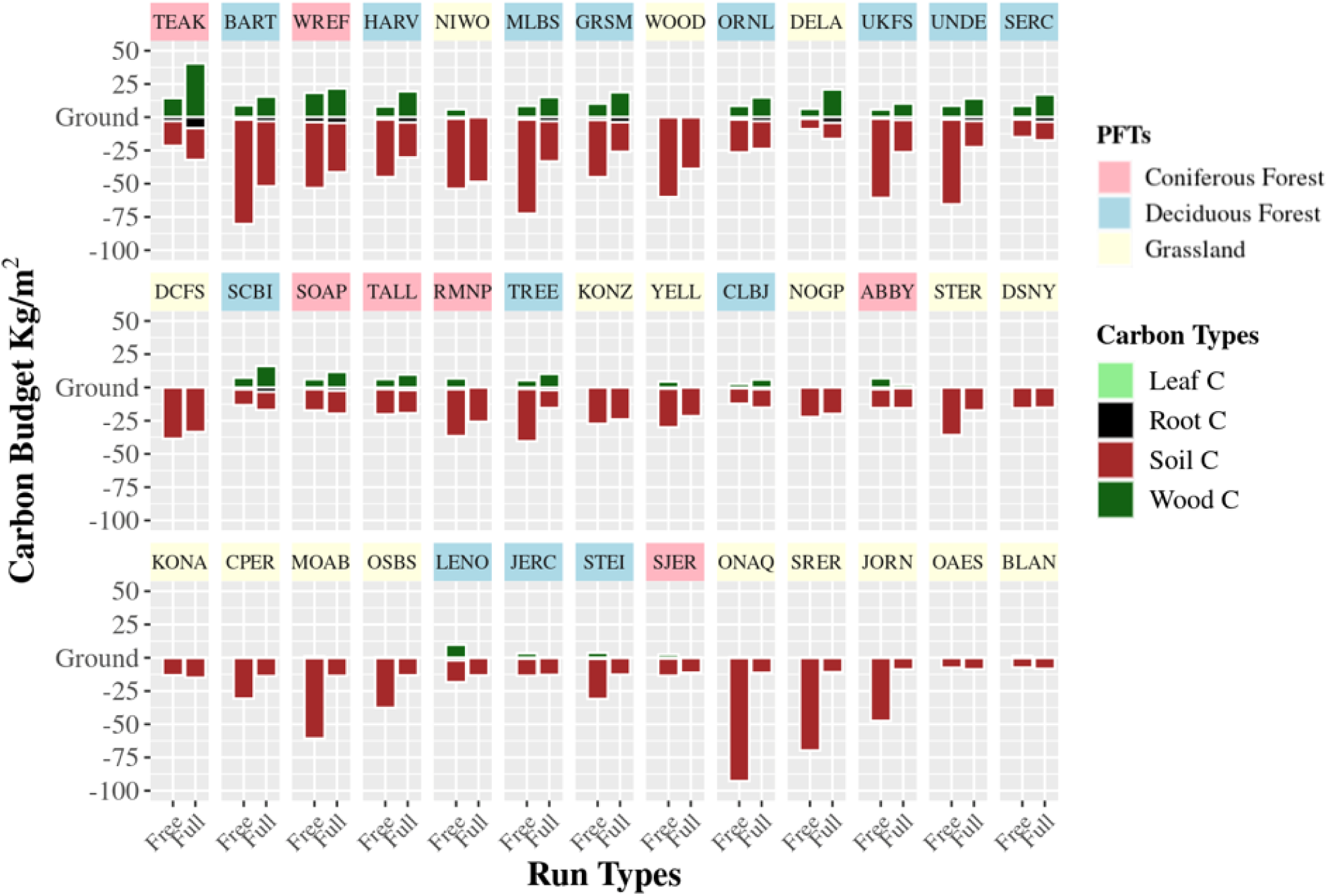
Carbon pools for the CONUS NEON Sites in 2021, sorted by total carbon stocks in the full assimilation. Each panel title is colored based on the corresponding plant functional types.

We also plotted the C pool uncertainties for each site (Figure 9) before and after the SDA based on equation 2 across four C budgets: leaf, wood, root, and soil C. Soil C dominates overall C uncertainty before (59.9 kg C/m^2^, 93.8% of total) and after (17.5 kg C/m^2^, 85.5% of total) the SDA experiment, especially across deciduous forests (28.0 kg C/m^2^, 93.4% of total) before and after the SDA experiments (9.7 kg C/m^2^, 85.1% of total) and grasslands before (39.3 kg C/m^2^, 92.8% of total) and after (13.0 kg C/m^2^, 88.2% of total) the SDA experiments. In contrast, for the coniferous forest, the uncertainties between wood and soil C are sometimes comparable (such as TEAK, SOAP, TALL, ABBY, and SJER) in the free run (4.0 kg C/m^2^, 56.9% of total) and the full run results (3.4 kg C/m^2^, 61.9% of total). Beyond that, the C uncertainties are reduced significantly (ANOVA stats: F(1, 349) = 71.3, P < 0.001 ***) by our SDA workflow, with an overall uncertainty reduction of 43.4 kg C/m^2^ (68.0% of total).

**Figure 9.**
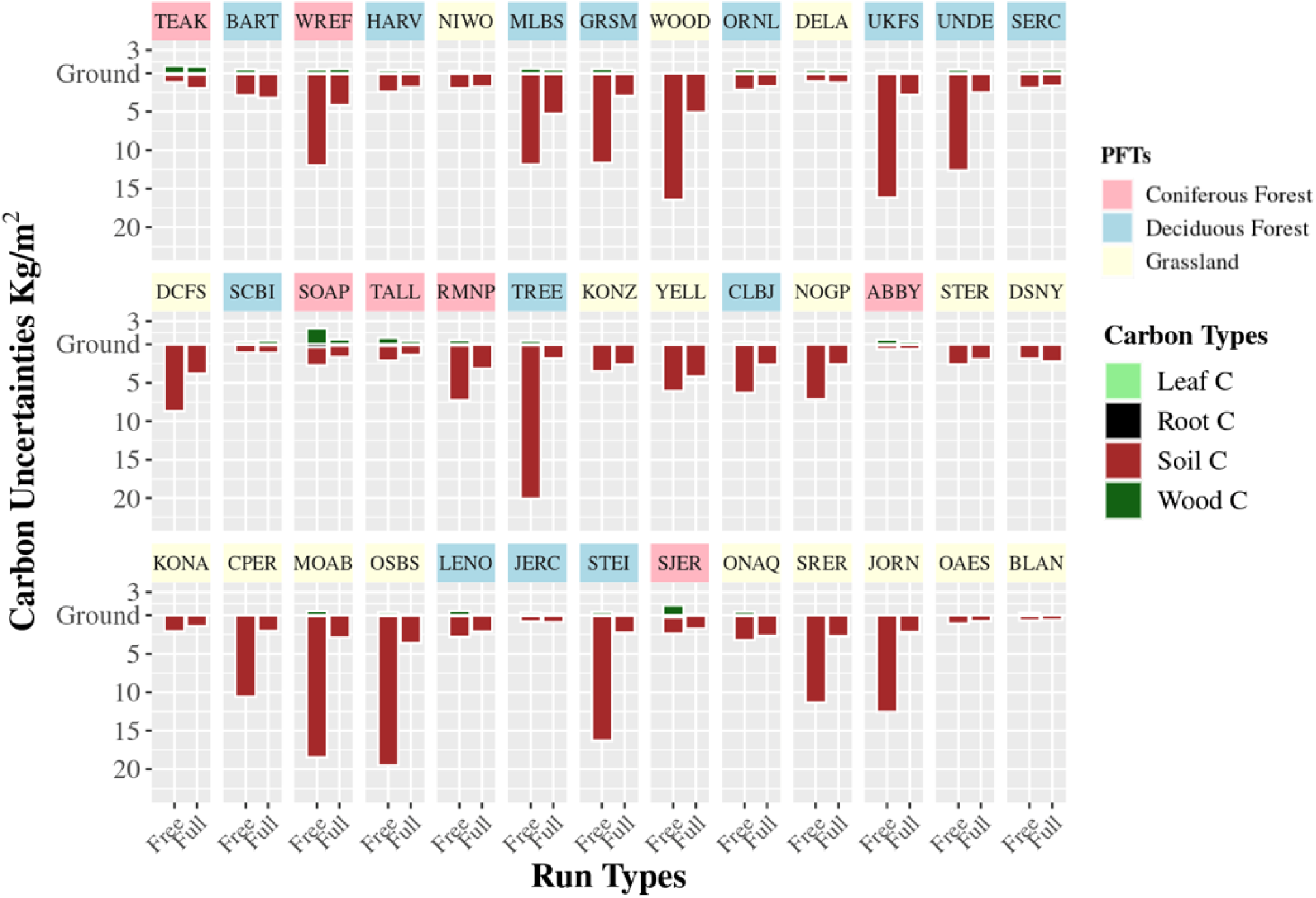
Uncertainties of carbon pools for the CONUS NEON Sites in 2021, sorted by their total carbon stocks. Each panel title is colored based on corresponding plant functional types.

Aggregating C uncertainties across PFTs we found that the “free run” C uncertainties were highest for deciduous forests (8.25 kg C/m^2^), followed by the grassland (7.35 kg C/m^2^) and conifers (5.02 kg C/m^2^) (supplemental Figure S1 A). The C uncertainties after SDA are evenly distributed across all three PFTs (2.67, 2.51, and 2.54 for the deciduous broad-leaf forests, grassland, and conifers, respectively), making the deciduous forest the highest C uncertainty reduction (67%), followed by the grassland (66%) and conifers (49%). Aggregated across different *level 1 EPA* ecoregions, C uncertainties are lowest in the Northwestern Forested Mountains, North American deserts, and Mediterranean California ecoregions, both before and after SDA (supplemental Figure S1 B). C uncertainties are largest, both before and after the SDA, across Northern Forests (6.76 kg C/m^2^ before and 1.87 after, 72% reduction), followed by Eastern Temperate Forests (5.02 kg C/m^2^ before, 1.29 after, 74% reduction) and the grasslands and croplands of theGreat Plains (2.8 kg C/m^2^ before, 1.49 after, 47% reduction).

Another factor hypothesized to affect C uncertainty is the size of the overall C stocks (e.g., sites with larger C stocks are expected to have higher C uncertainty). For the free run there is a positive linear relation (slope of 0.12) suggesting that larger C are associated with higher uncertainty (Figure 11), while in the full run the relationship is much more flat (slope of 0.03, Figure 11) indicates that the SDA reduces the sensitivity of C uncertainty to the C stock size.

**Figure 11.**
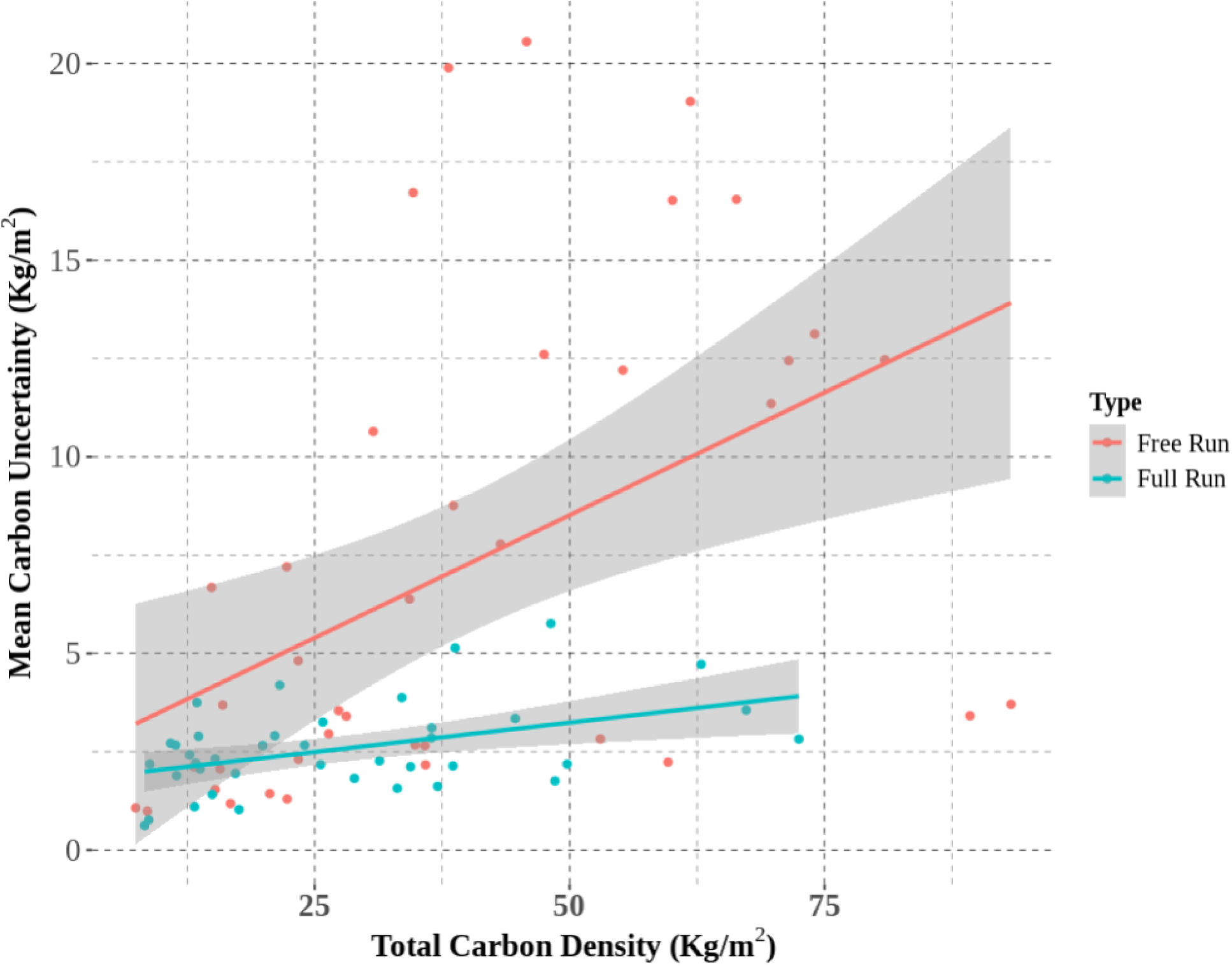
The relationship between total carbon stock density and carbon uncertainty for free run and 500km full run across NEON sites.

### 3.2. Across-variable VOI

VOI results were summarized into four panels (Figure 12), within which the upper/lower left blocks show the uncertainty increment of the four state variables (wood carbon, LAI, soil water, and soil carbon) relative to the full runs (top) and free runs (bottom), respectively. The actual values for each cell are reported in Table S1. The upper/lower right blocks represent the uncertainty increment of four key fluxes (GPP, transpiration, autotrophic respiration, and heterotrophic respiration), none of which were directly constrained by the SDA. When adding individual data constraints to the free run, we observed substantial direct constraints for the VOI of four state variables with uncertainty reductions of 39% (wood C), 75% (soil moisture), 88.3% (LAI), and 90.5% (soil C). Similarly, there was an increase in uncertainty when removing the corresponding observations from the full run, ranging from 36.1% (wood C) to 73.2% (soil moisture), 86.7% (LAI), and 91.8% (soil C).

**Figure 12.**
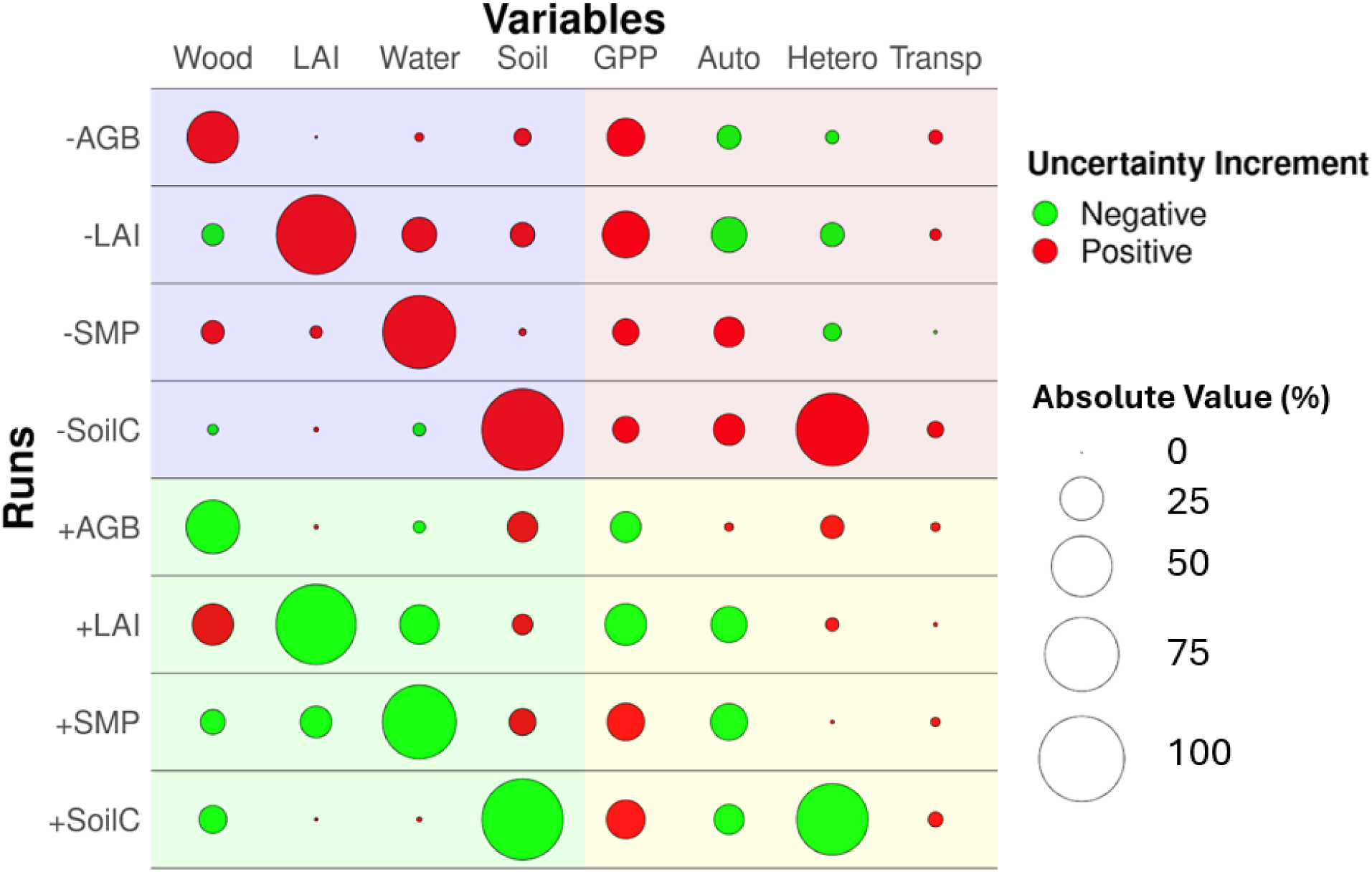
The across-site average VOI contribution of different runs to the uncertainty increment of water/carbon pools/fluxes. Here, we calculate the relative uncertainty increment between the plus-one observations run and the free run and between the minus-one observations run and the full run. We split the figure into four panels: the upper and lower panels show comparisons for free run and full run, while the left and right panels show the comparisons for the state variables and fluxes.

For the indirect constraints generated by across-variable correlations, we observed paired indirect constraints for the upper and lower panels among AGB, LAI, and soil water. For example, assimilating SMAP soil moisture led to uncertainty reductions of 8.1% and 13.3% for wood and leaf C pool estimations, while adding Landtrendr AGB and MODIS LAI led to soil moisture uncertainty reductions of 1.7% and 20.6%, respectively. The correlation between soil moisture and LAI is also supported by our relative change analysis (Figure 13A), which shows that increases in soil moisture within the SDA are associated with concomitant increases in LAI. For wood carbon, we counterintuitively observe that the assimilation of MODIS LAI increases uncertainty by 22%. Figure 13B, illustrates the relationship between changes in LAI and the uncertainties of wood carbon for the conifer forest PFT, such that the assimilation of MODIS LAI leads to an increase of LAI, which in turn leads to an increase in AGB, which leads to an increase in the absolute uncertainty of the woody biomass. For the interactions between wood and soil carbon pools, we observed decreases in wood carbon uncertainties by 10% when adding observations of SOC, while adding observations of AGB will lead to an uncertainty increment of 12% for SOC.

**Figure 13.**
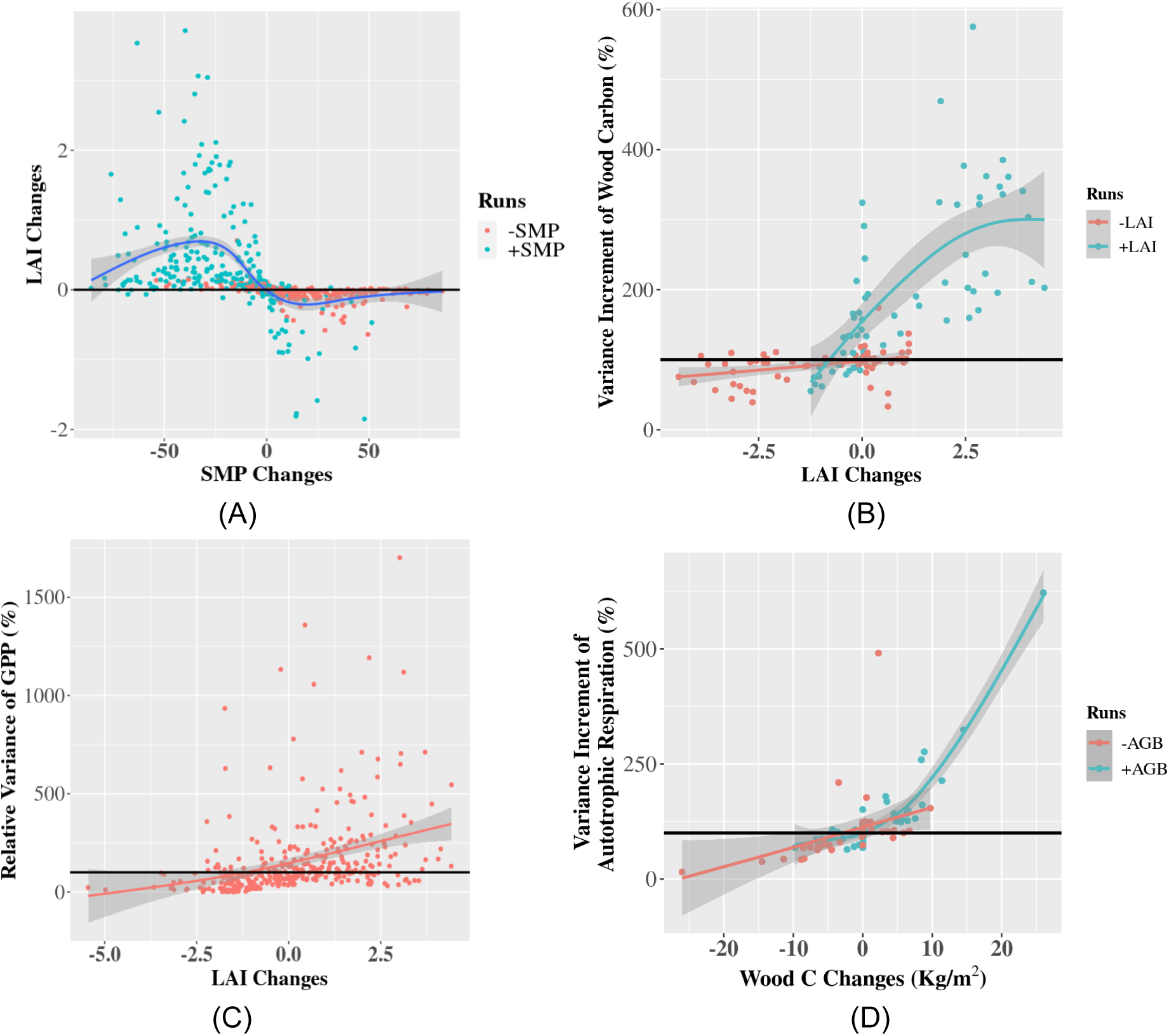
How changes in the size of one directly-assimilated pool are associated with the indirect changes in the relative uncertainties of another pool of flux. Color indicates whether the directly assimilated pool was added (blue) or removed (red) from different SDA runs (Table 2). Figures A and C represent the impact of soil moisture assimilation on the LAI changes and the uncertainty increment of GPP by the LAI changes. Figure B represents the uncertainty increment of woody biomass over coniferous forest sites when assimilating observations of MODIS LAI, while Figure D represents the uncertainty increment of autotrophic respiration when assimilating observations of AGB.

For the VOI of the four fluxes (right-hand blocks in Figure 12), assimilating SOC reduced the uncertainty of heterotrophic respiration by 70.5%, while integrating both MODIS LAI and Landtrendr AGB decreases the uncertainty of GPP by 29.7% and 19.1%, respectively, because LAI is a critical variable in the prediction of GPP within the SIPNET model calculation (Figure 13C). By contrast, assimilating soil moisture leads to an increase in the GPP uncertainty by 19%, which may be associated with the strong correlation between soil moisture and LAI (Figure 13A) and the previously mentioned correlation between LAI and GPP uncertainty (Figure 13C). Regarding respiration fluxes, assimilating soil moisture and SOC decreased the uncertainty in autotrophic respiration. By contrast, the assimilation of LAI, AGB, and soil moisture tended to increase the uncertainties of heterotrophic respiration. We also observed that adding the Landtrendr AGB observations increased the uncertainty of autotrophic respiration by 1%, while the uncertainty decreased by 7.3% when removing AGB observations. Part of this response is related to the increase in wood C within the full run (Figure 8), as Figure 13D shows that increasing wood carbon is correlated with an increase in autotrophic respiration uncertainty.

Aggregating the VOI analysis to the level of the overall C stock assessment (see Figure 14, where we calculated the standard error 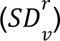 of each C pool using the same eq 3), SOC dominates the overall C uncertainties in all runs (Table 2), wood C is consistently the second largest uncertainty, and leaf C uncertainties are substantially smaller. That said, the assimilation of SOC within the SDA is also the most valuable data constraint, leading to the single largest total C uncertainty reduction (65.5%). Counterintuitively, we also observed that adding observations of AGB, LAI, or soil moisture to the free run without also assimilating SOC led to increases in the total C uncertainty because adding these observations increased soil C uncertainty (Figure 12) more than it decreased the uncertainty in each of these pools. Assimilating all observations will result in the lowest total uncertainty because of the direct and indirect across-variable benefits discussed previously.

**Figure 14.**
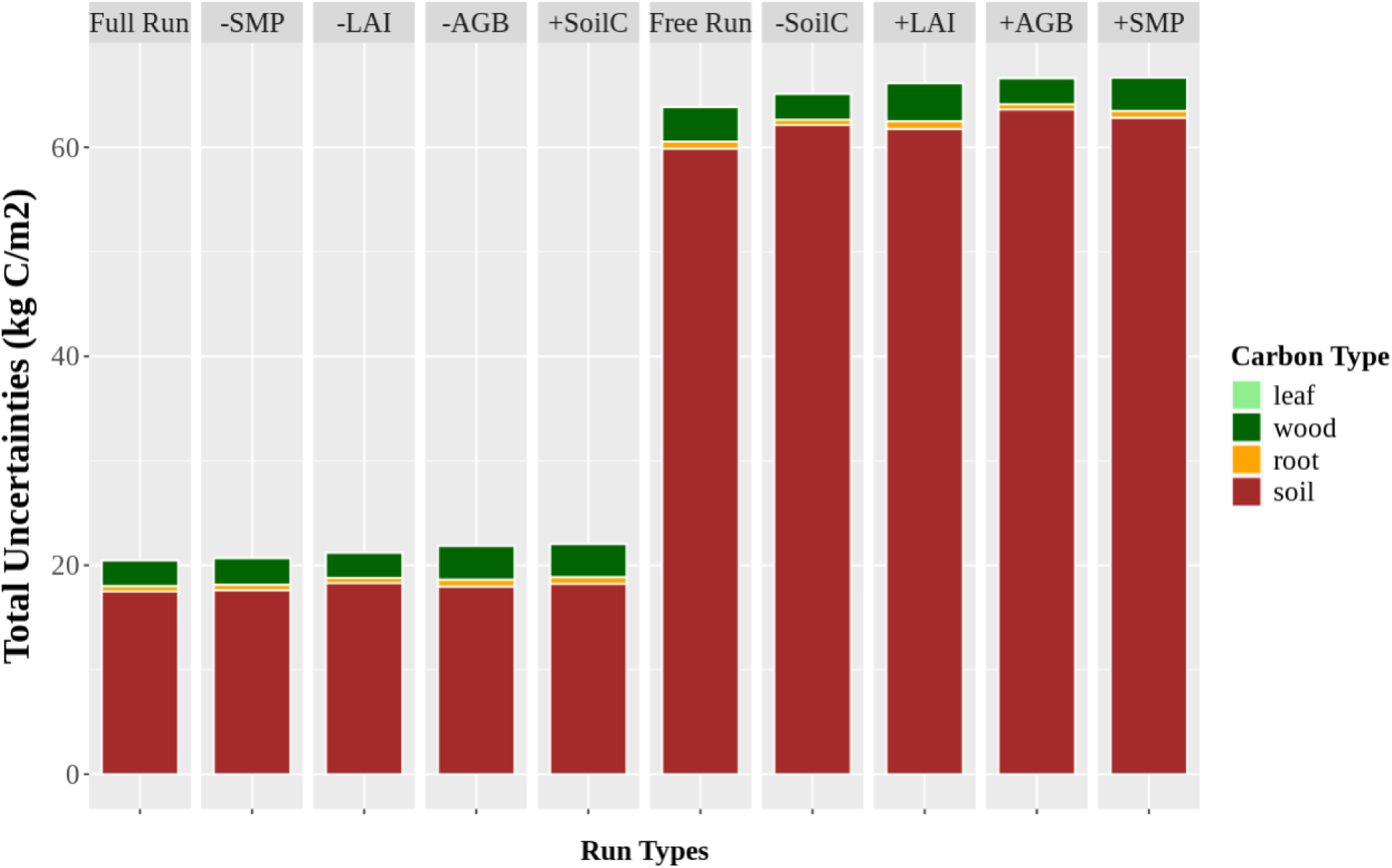
Uncertainties of the overall carbon stock (including leaf, wood, roots, and SOC) for different SDA runs (see Table 2) sorted in order from smallest to largest total uncertainty.

### 3.4 Across-space VOI

The SDA framework incorporates information from nearby locations as a function of how strongly model ensemble members are correlated across different locations, rather than simply as a function of distance. However, we can assess how VOI changes across-space by varying the spatial scale of localization (i.e., the search window over which correlations are calculated). Figure 18A shows the trends of our four state variables when we increase the localization spatial scale from 0 to 500 km with a 100km interval, expressed in terms of further uncertainty reduction relative to the “full run” with 0 localization. Overall, LAI has the largest and the most consistent uncertainty reduction (∼35%) across spatial scales, while wood carbon has the second largest uncertainty reduction (∼30%), though with some sampling variability (at 100 and 400 km) within the trend. SMAP and SOC share a relatively slow but more stable trend (with ∼ a 15% reduction at 500km) across space.

**Figure 18.**
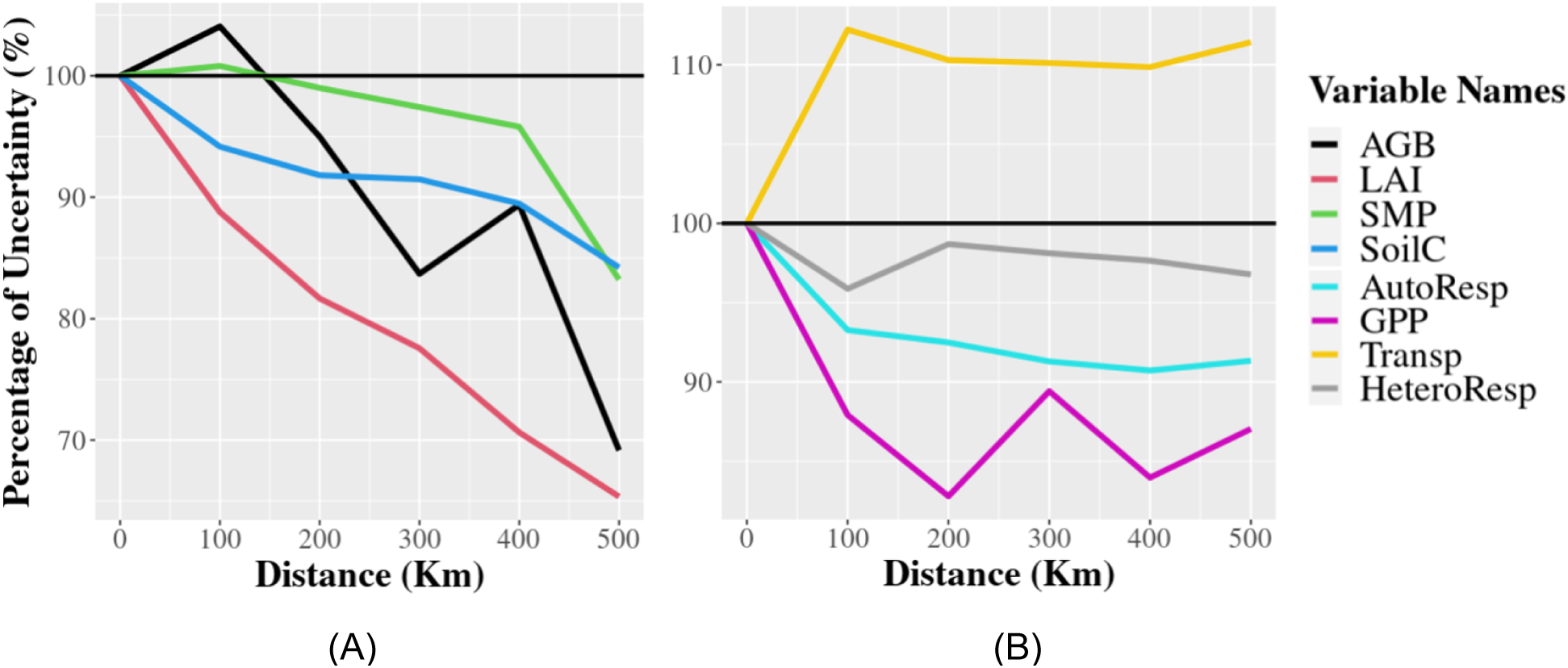
The overall trend of uncertainty changes of (A) state variables and (B) fluxes across different spatial scales for four state variables and fluxes. Here, we standardize the 100% baseline (horizontal black line) as the uncertainty of state variables or fluxes at 0 km, and any points after that will be calculated against the baseline.

Figure 18B shows the spatial trends for the corresponding C and water fluxes. The largest changes in uncertainty occur when going from 0 km to a 100 km scale, followed by roughly flat trends suggesting no substantial changes in flux uncertainties at larger distances. GPP benefits the most from shared spatial information, reaching its minimum uncertainty at 200 km (17% reduction). Autotrophic respiration had the second largest constraint (9% at 400km), while heterotrophic respiration only saw a 3% reduction (at 100km), and the uncertainty of transpiration increased by >10%.

Figure 19 shows the spatial pattern across CONUS of how uncertainty changed as a function of localization distance (i.e., whether or not sharing information spatially was beneficial [green] or not [red]). For the wood carbon map, most grassland sites showed no response to the AGB assimilation (flat trend) because wood carbon is absent from these systems. The southern California sites (TEAK, SJER, and SOAP) had highly variable amounts of wood carbon, which might explain why sharing spatial information increased uncertainty at the TEAK site. Moreover, the HARV and BART sites also share conflicting information, increasing the uncertainty of AGB for the HARV site. Both southern California (TEAK, SJER, and SOAP) and northeastern site groups (HARV and BART) are spatially clustered while having variabilities in wood C budgets, which contribute to the increase of AGB uncertainty at the 100 km scale in Figure 17A.

**Figure 19.**
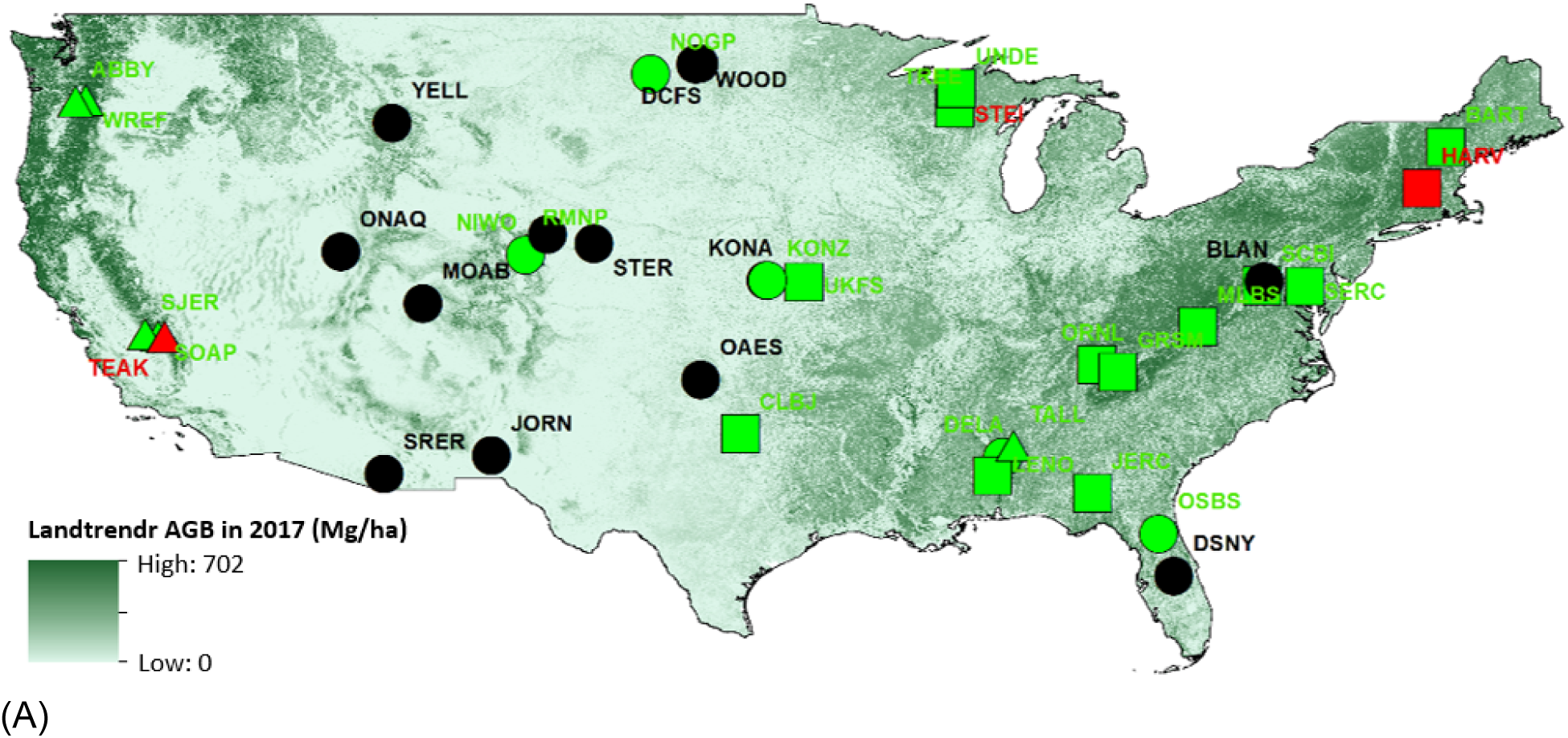
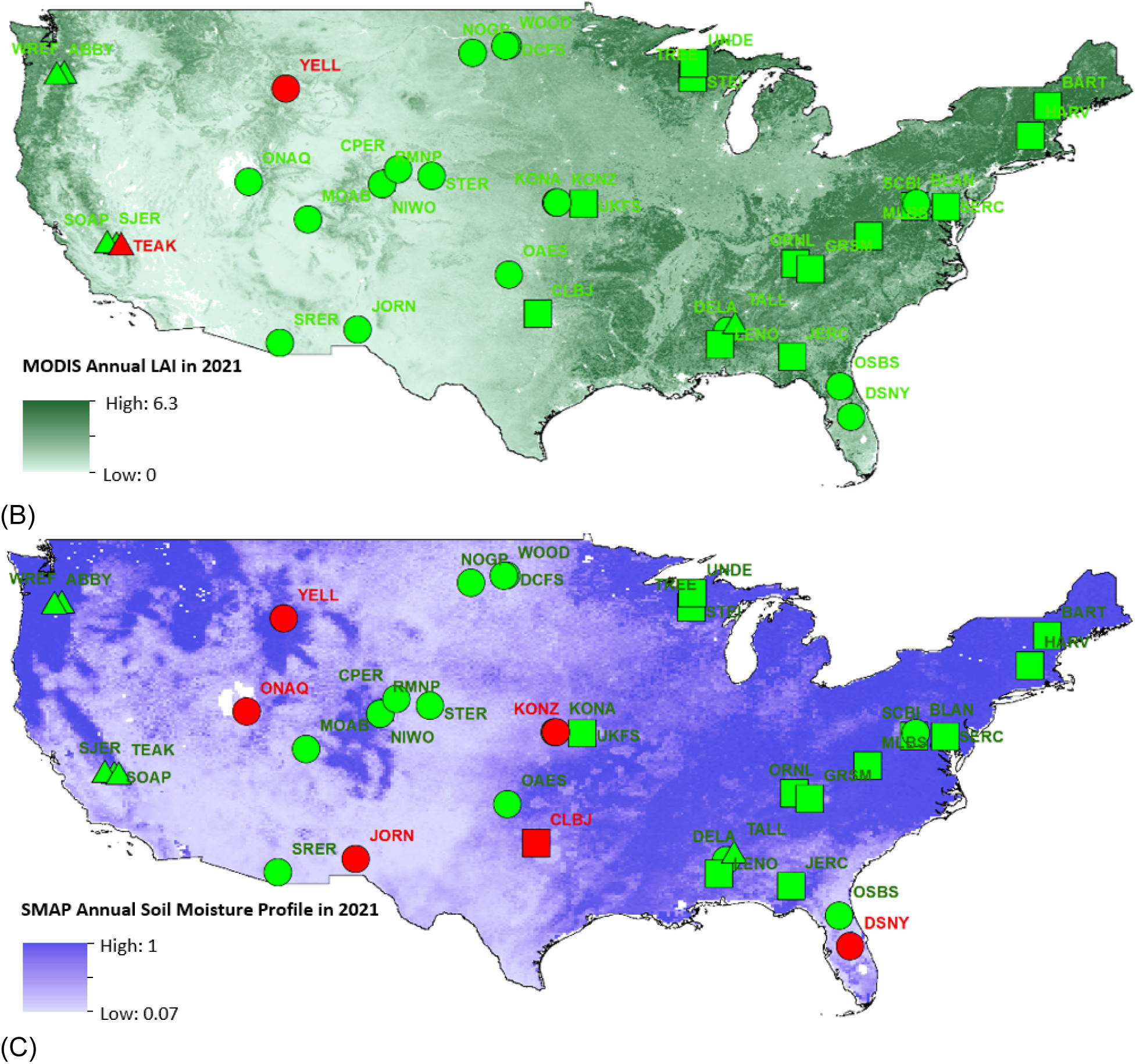
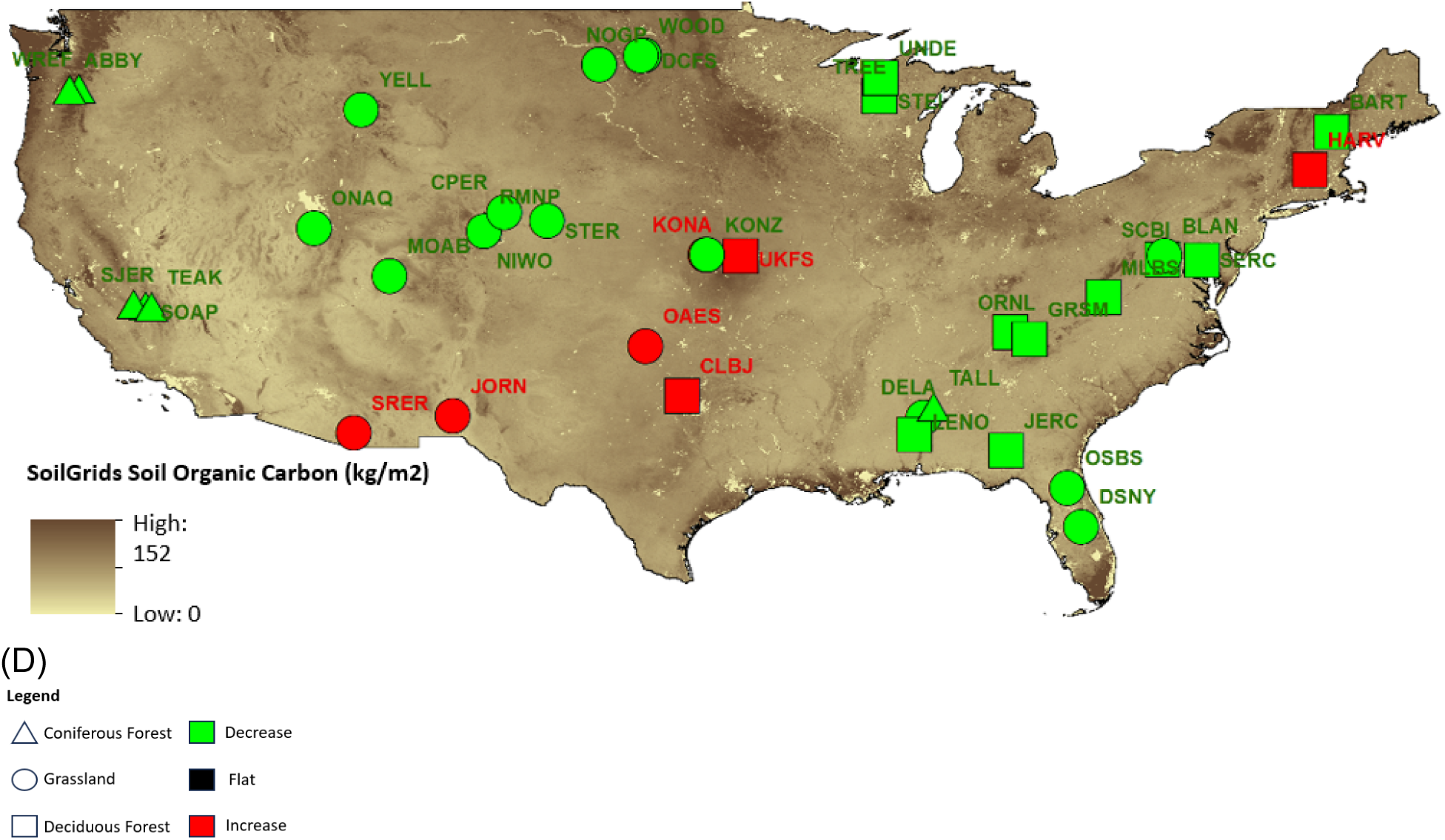
The spatial patterns of the spatial value-of-information across different scales. Colors indicate whether uncertainties increase, decrease, or remain flat as the spatial scale of localization is increased. We finally plotted the classified slopes along with the corresponding plant functional types of each location, with the background of each observation. Figures A-D represent the spatial patterns of AGB, LAI, SMP, and SOC, respectively.

The LAI map showed reduced uncertainty as a function of localization range across almost every site in the CONUS scale except for the YELL and TEAK sites, leading to the most consistent decreasing trend on average (Figure 18A, red line). In contrast, more sites share conflicting information spatially for the SMP and SOC variables. As we can see from Figure 19C, the NEON sites within eastern and western US regions are generally well-constrained, while in the mid-US, where the SMP varies dramatically spatially, as suggested by the background annual SMAP map, the spatial information will increase the uncertainties among YELL, QNAQ, JORN, CLBJ, and KONZ sites. Unlike SMP, the SOC map (Figure 18D) shows generally well-constrained outcomes by sharing information spatially, except for the central (KONA, UKFS, OAES, and CLBJ) and southwestern (SRER and JORN) US, where the red-colored NEON sites are clustered.

### 3.5. Cross-validation: Across-Variable Residual Errors

In addition to the across-variable uncertainties, the VOI experimental design (Table 2) allows us to cross-validate the SDA data product’s residual errors, 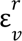, which are summarized in terms of the MAE between the held-out observations and the posterior mean of the reanalysis results for variable *v* and experiment *r*, averaged across time and space. Table 3 shows that assimilating “indirect” observations had little impact on the residual errors for wood carbon (from 4.68 to 4.66kg/m^2^), LAI (from 1.33 to 1.57), soil water (from 26.49% to 26.29%), and SOC (from 21.3 to 21.2kg/m^2^) when comparing the free run to the experiment that excluded that specific variable from the full run. The residual errors for assimilating the direct observations showed significant improvements compared to the free run results: wood carbon (from 4.68 to 0.34kg/m^2^), LAI (from 1.33 to 0.92), soil water (from 26.49% to 1.50%), and SOC (from 21.3 to 4.8kg/m^2^).

**Table 3.** The MAE of C and water budgets under different experiments (see Table 2). The diagonals of adding or subtracting the corresponding observations are bolded below. The bar plot also labels the relative magnitude for each column. Here, we summarize the MAE of AGB, LAI, SMP, SOC, and the combined AGB+SOC+Leaf to show the impacts of SDA on the residual error across C and water budgets.

| Experiment Name | AGB (kg/m<sup>2</sup>) | LAI (m<sup>2</sup>/m<sup>2</sup>) | Soil Moisture (%) | SOC (kg/m<sup>2</sup>) | Total C MAE (kg/m<sup>2</sup>) |
| --- | --- | --- | --- | --- | --- |
| Full Run | 0.300 | 0.940 | 1.452 | 4.805 | 5.105 |
| -AGB | **4.659** | 0.940 | 1.441 | 4.632 | 9.291 |
| -LAI | 0.305 | **1.574** | 1.432 | 4.791 | 5.096 |
| -SMP | 0.344 | 0.915 | **26.293** | 4.553 | 4.897 |
| -SoilC | 0.313 | 0.942 | 1.447 | **21.196** | 21.509 |
| +AGB | **0.339** | 1.357 | 26.564 | 21.845 | 22.184 |
| +LAI | 4.664 | **0.924** | 26.276 | 20.651 | 25.315 |
| +SMP | 4.635 | 1.536 | **1.499** | 21.382 | 26.017 |
| +SoilC | 4.660 | 1.325 | 26.596 | **4.822** | 9.482 |
| Free Run | 4.675 | 1.333 | 26.493 | 21.255 | 25.930 |

The total C residual error column shows that SOC dominates the patterns of the total C residual error. Furthermore, the SOC and total C residual errors reach the lowest point (4.553 and 4.897, respectively) for the run that excludes SMAP, not the full run, while the second highest SOC and the highest total C residual errors (21.382 and 26.017, respectively) occur when we only assimilate SMAP observations, rather than the free run. Thus, while assimilating SMAP reduced uncertainties about soil moisture, it may have a small negative impact on uncertainties in total C stock.

## 4. Discussion

In this study, we employed an ensemble-based Bayesian SDA approach across CONUS NEON sites that simultaneously assimilated four observations (LAI, AGB, Soil moisture, SOC) from 2012 to 2021. In addition to providing an essential proof-of-concept for the ability to simultaneously assimilate a large number of data constraints to constrain terrestrial C pools and fluxes, this work was motivated by five key research questions (sec 1.1), each of which is addressed in a separate subsection below.

### 4.1 How well did we assimilate the four major C pools into TBMs? What are the major impacts of data assimilation on the general C budgets?

Our SDA framework efficiently assimilates data on four major C and water pools into a terrestrial biosphere model, reducing uncertainty across different variables and spatial domains from 39 to 90.5%, and reducing overall C budget uncertainty by an average of 65.5%. This reduction in predictive uncertainty is also consistent with a corresponding reduction in residual error compared to the free-run results. Reducing C budget uncertainties help us better understand the spatial-temporal states and transitions within the C cycle, benefiting both basic science and MRV efforts. Beyond that, more accurate estimations can help maximize the efficiency of limited monitoring resources and improve our ability to mitigate carbon loss by increasing confidence about which stocks are in decline.

In addition, comparing C cycling reanalysis results to “free run” model simulations produced estimates that, on average, contain more wood and less SOC (Figure 8). By reconciling model simulations with real-world observations, the iteratively updated model simulations become both more precise (i.e. less biased) and more accurate over time.

After SDA, C uncertainties were also relatively consistent across PFTs, though they did vary by ecoregion (Figure 10B), suggesting differences in model performance across plant species and space. Efforts are underway to build upon the existing hierarchical Bayesian calibration of SIPNET (Fer et al., 2018, 2021), aiming to further improve model calibration and better capture spatial heterogeneity in optimal model parameters.

While the SDA was successful overall in reconciling SIPNET with multiple data constraints, there are places where the current SDA system could be improved in the future. A key one is in better reconciling the spatial-temporal extents between different data sources (model, observations, ICs, etc). For example, our analyses showed a better correspondence between NEON AGB initial conditions and the 30m LandTrendr AGB when the latter was aggregated to larger spatial scales (Figure SX). This has not yet been incorporated into the SDA as existing biomass data products are missing estimates of spatial autocorrelation and so additional research is required to determine how to aggregate the uncertainties in this product accurately (Kennedy et al 2024, Wadoux 2023). Beyond that, the Landtrendr AGB also has limitations on both temporal and spatial scales (e.g., only available within CONUS U.S. and until 2017). The opposite challenge exists with SMAP, which has a fairly coarse resolution (9 km) and a relatively late start date (2015), such that estimates might be improved if means and uncertainties were disaggregated, though this is lower priority given the small contribution of SMAP to overall uncertainty budgets. Much more urgent is SOC, which was the dominant uncertainty but only provided one snapshot for each pixel rather than a time series (see section 4.2).

In the future, spatial aggregation and downscale techniques could be used to standardize the spatial resolution across data sources. For example, Xu et al. (2022) downscaled SMAP SM to 1km, while Rodríguez-Veiga et al. (2024) aggregated AGB estimations from Lidar and Spectral images. Better harmonizing spatial scales may also require explicitly accounting for the sub-grid fractional distribution of PFTs (Hartley et al., 2017), which could be achieved by aggregating higher-resolution remotely sensed land cover maps (e.g., Sentinel-2) to better represent plant composites across the landscape. There are also analogous opportunities to improve the temporal resolution. For example, an improved seasonally-to-monthly SDA frequency is likely to improve the information contribution for variables with shorter memory (e.g., LAI and SM).

A crucial next step in this work is to scale up to larger spatial extents to be able to both meet the demands for improved C MRV and to use the reanalysis technique to improve our understanding of the interactions between C cycling and environmental and climatic conditions (Migliavacca et al., 2009). While our SDA framework has demonstrated the capacity to assimilate numerous observations simultaneously, providing comprehensive uncertainty quantification and propagation and reducing uncertainties across the CONUS NEON scale, transitioning to larger spatial domains presents several technical challenges: 1) The localization approach used in our study, which depends on distances among locations, should be improved to consider variations in ecoclimatic and topographical properties, not just physical distance (Serra et al., 2011); 2) The computational efficiency of the current workflow might become a bottleneck for the large-scale analysis, and therefore we might consider using hybrid methods such as machine learning-based SDA approaches (Farchi et al., 2021); 3) With increasing heterogeneity and dimensionality, there is a growing need for more comprehensive uncertainty quantification and propagation. While our current workflow employs Gamma-Normal sampling methods, which may ignore covariance between process errors, future considerations should involve utilizing the Wishart-Multivariate Normal sampling method to quantify and constrain process errors jointly (Dokoohaki et al., 2022).

### 4.2 What data sources have the greatest impact on constraining the C cycle directly?

As indicated by our C uncertainty analysis (Figures 9 & 14), the assimilation of SOC data had the largest impact on constraining the C cycle directly, with a decrease of 41.8 kgC/m² between the free and full runs, resulting in a 90.5% reduction in SOC uncertainty and a total carbon uncertainty reduction of 65.5%. Despite this, SOC still dominates the overall C budget uncertainties, both before and after the SDA. Qianyu et al. (2024, in review) provided a more detailed analysis of how the SoilGrids SOC observations interacted with the SOC prediction and the indirect impacts of SOC brought by factors such as observations from the rest of the C pools, time of SDA, and spatial indirect correlations.

Further reductions in SOC uncertainty are clearly a top priority, but there are challenges and opportunities facing an improved assimilation of SOC. For example, SOC is difficult to model at a large scale because it varies significantly spatially and is heavily dependent on the physical properties of soil, as well as climatic and environmental conditions (Casson et al., 2024). Additionally, SOC is complex to measure due to its variability in space and depth. Remote sensing observations provide a means to correlate SOC estimations with multi-to hyperspectral observations (Angelopoulou et al., 2019; Castaldi et al., 2019). However, assimilating such products needs to be done carefully so as to not double-count the indirect constraints already coming from the assimilation of existing multi-spectral observations, such as MODIS LAI and Landtrendr AGB, nor double counting that the SOC data used to construct such products may not be independent of the SOC being assimilated. Similarly, while different SOC databases and maps exist (SoilGrids, Poggio et al., 2021; Soil Survey Geographic Database (SSURGO), Sundquist et al., 2009; GSOCmap V1.6, FAO, 2022; the international soil carbon network (ISCN) Nave et al., 2016; etc.) large scale SOC maps contain substantial amounts of duplicate the information (i.e., they are often using the same underlying field measurements), making the harmonization of existing data products difficult, if not impossible. In an ideal world, the lack of independence of both ground-based and remotely-sensed products could be accounted for via observation error covariance matrices, but the post-hoc construction of such estimates would be a substantial undertaking in its own right. Last but not least, different SOC databases and maps provide measurements to different depths (e.g., 30cm vs. 200cm), and these depths may not match the depths in the model, which can make it challenging to align observations with what a model is predicting (e.g., SIPNET does not use layered soil biogeochemistry).

In addition to reducing SOC uncertainty, our next step is to improve the modeling and assimilation of AGB, as it is the second-highest pool and source of uncertainty among the overall C stocks and a key component in C cycling that relates to both above- and below-ground biomass. As we discussed earlier, the AGB uncertainties and residuals may arise from discrepancies in resolution, such that the uncertainties may increase when datasets with varying resolutions are combined. To resolve this, we will need to consider the spatial aggregation techniques in reconciling spatial representations between datasets. The spatial autocorrelations for the uncertainty maps should be examined and modeled carefully, as the aggregation is highly dependent on this process (Wadoux & Heuvelink, 2023).

Beyond that, this study also shows an increased uncertainty of SOC (Figure 14, column +AGB) when assimilating LandTrendr AGB. Therefore, we will also need to reconcile the different proxies of C cycling in the DA and modeling by matching resolutions and physical mechanisms between datasets and modeling processes, such that assimilating AGB will improve performance with reduced uncertainty and a smaller residual in other pools. Additionally, optical remote sensing may be saturated when attempting to measure dense forests, rendering the predictions unreliable and, similarly, our DA results. Therefore, our SDA workflow should incorporate other data sources such as LiDAR platforms (e.g., ICESat-2 and GEDI) and forest inventory data.

### 4.3 Which indirect constraints (i.e., based on model covariances) are strongest across pools and between pools and fluxes?

This study demonstrates that C budgets can be substantially constrained indirectly by sharing information across observations, ecological processes, and spatial scales. For instance, because of the correlations between soil moisture, leaf area index (LAI), and aboveground biomass, assimilating SMAP soil moisture lead to uncertainty reductions of about 8.1% and 13.3% for wood and leaf C pool estimations while adding Landtrendr AGB and MODIS LAI lead to soil moisture uncertainty reductions of 1.7% and 20.6%, respectively. It is also essential to consider the relative VOI changes and the absolute C uncertainties jointly. For example, because SOC dominates the overall C uncertainties (Figures 9, 12, and 13), the indirect constraints associated with assimilating AGB, LAI, or SMP (12.3%, 5.5%, and 9.6%, respectively) lead to nontrivial increments for the SOC absolute uncertainties (3.7, 1.9, and 3.0 kg C/m2, respectively). Overall, such indirect constraints allow us to reduce uncertainties about ecosystem variables that may be challenging to constrain directly due to the limited availability of measurements or high observation uncertainties. More broadly, the feature of sharing information across variables, ecological processes, and space facilitates is not just a mechanism for reducing uncertainties but also represents a successful reconciling of different sources of information, representing areas of agreement among different types of data and between data and model.

Additionally, indirect interactions lead to significant uncertainty reductions for otherwise unconstrained variables. In this study, we were able to use indirect constraints to reduce uncertainties about ecosystem C fluxes, for example constraining gross primary productivity (GPP) when assimilating MODIS LAI and autotrophic and heterotrophic respirations when assimilating SoilGrids SOC. While the ability to constrain fluxes indirectly based on the assimilation of data about pools is a valuable feature of SDA, an important future direction to further reduce these uncertainties would be to assimilate flux data directly, such as eddy covariance observations from networks like NEON, Ameriflux, and Fluxnet (Keller et al., 2008; Novick et al., 2018; Baldocchi et al., 2001). An essential opportunity for land DA is that flux posteriors can serve as priors for atmospheric inversion modeling (Peylin et al., 2013), helping to reconcile top-down information about the carbon cycle from the atmosphere with bottom-up information from in situ and remotely sensed data about the land. Such an approach would provide a harmonized understanding of land-atmosphere interactions, ultimately benefiting the goal of a comprehensive Earth-surface SDA system.

### 4.4 Which indirect constraints are strongest across space, and how quickly does information sharing drop off with distance?

The spatial VOI analysis demonstrates that the uncertainties of all four pools, and three of the four fluxes analyzed (all except transpiration), decrease as the spatial scale of information sharing increases from 0 to 500 km. Specifically, LAI and GPP are the most constrained C pool and flux, respectively. This is likely because LAI estimates are more comparable across space and PFTs, which will likely lead to relatively higher spatial correlations for LAI, and GPP benefits from a strong indirect constraint from LAI (Sec 4.3). In particular, because in our examples, there are no covariances in the observation or process errors, indirect constraints come from covariances across ensemble members generated by their shared responses to their shared ensemble inputs (e.g., parameters, meteorological drivers, initial conditions), and the partitioning of the relative contributions of each input to such shared spatial information would be a useful future direction. Indeed, when we examine pools across various spatial scales, all appear to continue declining in uncertainty, rather than asymptotically, even at distances of 500 km, suggesting substantial spatial coupling across ensemble members in model outputs. While we are not aware of other papers that have similarly leveraged ensemble covariances to compare to, such spatial scales are larger than we initially expected, though not inconsistent with the literature on ecological teleconnections (Garcia et al., 2016).

Looking beyond LAI, because of the limited number of sites in this study, the spatial sharing of information will also be limited, and sometimes even in conflict, if there is no correlation between nearby sites. Such patterns help explain the patterns of shared spatial information for other pools. For example, the AGB uncertainty increases at the 100 km scale (Figure 18A) because some clustered sites (e.g., the southern California sites, TEAK, SJER, and SOAP) have highly variable amounts of wood carbon, which contributes to the increased conflicts when they are grouped. The SOC and SM are relatively less constrained due to the spatial heterogeneities that exist in the central US (see the background map in Figures 19 C and D). As a whole, we thus observe that aboveground variables tended to benefit more from indirect spatial constraints than belowground variables, which suggests a stronger role of spatial covariances for vegetation (e.g., parameter covariances or shared responses to atmospheric variability) than for equivalent responses in soil biogeochemical and hydrologic processes. While it is entirely possible that vegetation ecophysiological processes are coupled over larger scales than soil processes (e.g., pedogenisis), especially given the high spatial heterogeneity of soils and the short lifespans of soil microbes, we are hesitant to read too much into the latter response as the representation of soil processes in SIPNET are fairly simple and do not explicitly account for the known spatiotemporal predictability of the soil microbiome (CITE Colin, Zoey) or the spatial patterns of many important soil covariates, such as pH and nutrients (CITE). Likewise, PFT-based approaches to the calibration of vegetation properties, such as we employ here, underestimate spatial parameter heterogeneity (Fer et al., 2021) and thus may contribute to an overestimation of aboveground spatial covariances. In addition, because the Tobit-Gamma Filter employed here does not account for spatial covariances in the model process error this could also be affecting the estimated spatial ranges of shared information, likely leading to an underestimation over shorter spatial scales as spatial error covariances tend to be positive. That said, it is unknown whether this would affect the overall spatial range of correlation, as spatial error covariances often decay rapidly with distance and can even become negative at larger spatial scales. Regardless, shifting back to a Tobit-Wishart Filter would allow for the estimation of such spatial error covariances, though any potential information gained would need to be balanced against the higher computational costs.

The spatial VOI patterns for the fluxes (Figure 18B) indicate that GPP is the most constrained variable, followed by autotrophic and heterotrophic respirations and transpiration with increased uncertainty compared to the baseline (at the 0 km scale). Unlike patterns for the pools, for which uncertainties are continuously reduced (although SMP and SOC exhibit a slower trend through space), there is little additional shared spatial information about fluxes beyond a 100 km scale, suggesting that fluxes are more indirectly related to the observations and the correlations within them. That uncertainty in transpiration increased when borrowing strength across space is slightly surprising but is not inconsistent with the within-site results, where transpiration showed the weakest indirect constraints among all the variables considered (Figure 12).

The ability to borrow strength spatially brings valuable opportunities to harmonize differences between “wall-to-wall” datasets (e.g., airborne or space-borne observations) and more sparse ground-based networks (e.g., NEON, FIA, LTER, etc.). For example, data from ground networks can serve as valuable “anchor sites” that tie down pools and fluxes with high-quality field data and share that information with neighboring sites, which would otherwise only be constrained by remote sensing. By identifying and mapping the impacts of shared spatial information on different times, locations, PFTs, and variables, and assessing the localization distance decay associated with such observations, we can better identify gaps in such networks and optimize monitoring programs. Valler et al. (2019) also point out the need to better adjust the localization scales to represent meaningful ecological continuity. For some highly spatially varied variables (e.g., C and water fluxes), assimilating observations with spatial covariance yields more accurate and less uncertain modeled outputs compared to those generated by spatially independent measurements (Ran et al., 2016). Furthermore, as suggested by our spatial-VOI analysis, when ecosystems are spatially heterogeneous, there is the potential to share conflicted information (Figure 19). In such cases, a traditional localization based on physical distance could be improved upon by using distances in environmental space, potentially weighting different ecoclimatic and edaphic variables by estimates of the sensitivity of model outputs to these different inputs. Finally, because increasing localization distances also increases the computational complexity resulting from the increased dimensionality of spatial covariances, these distances must be set to balance the computational demand required by these operations and the increased accuracy that sharing spatial information provides.

### 4.5 Are there places where different information sources conflict?

Within the VOI analysis, we identified a number of places where there were apparent conflicts among information sources. For example, the small decrease in AGB uncertainty (X%) associated with removing MODIS LAI observations from the assimilation suggests a small conflict among SIPNET, LandTrendr AGB, and MODIS LAI around the joint estimation of AGB and LAI. One possible explanation is the spatial mismatch in both mean and uncertainty between MODIS (500 m) and LandTrendr (30 m) observations. Beyond that, when assimilating MODIS LAI observations, the uncertainty increment of AGB (Figure 12) and the increased wood C for coniferous forest (Figure 13B) suggests that the conflicts between LAI observations and AGB estimations may also be related to differences between NEON AGB IC and MODIS LAI or conflicts in the parameterizations in the model. Related to this is the small negative impacts of assimilating AGB and LAI on the uncertainties in both autotrophic and heterotrophic respiration. For example, if the model underestimates the AGB forest, then “nudging” the trees larger will increase not only wood C, but also leaf C, Ra, and litter, and thus also Rh. The increase in leaf C may be in conflict with LAI observations, while the increase in pool sizes may increase absolute pool uncertainties even if the borrowing of strength reduces the relative uncertainties. Such conflicts may not have been detected in a traditional stand-alone calibration, where modeled state variables are not required to be in their “true” states when calibrating parameters, and thus nudging pools could introduce errors in fluxes. A joint estimation of model states and parameters with the SDA system may help reduce these sources of conflict associated with model parameter errors.

In addition to places where indirect constraints needed to be reconciled within the SDA, we also identified instances where the assimilated observations required reconciliation with the initial conditions. For example, the NEON SOC initial conditions were systematically higher compared to the SoilGrids SOC observations (see Figure 6), which resulted in decreased SOC estimations across the NEON scale after assimilating the SoilGrids SOC data. Similarly, AGB in our model simulations increased for forest ecosystems, but declined in predominantly non-woody systems, when Landtrendr AGB was assimilated. The latter, at least, is mostly due to the failure of the current model-data assimilation to capture heterogeneity within the NEON sites, as pixels that the assimilation classified as non-woody often had a sufficient number of woody patches at the NEON site scale for the NEON vegetation surveys to include forest inventory plots. Beyond that, our analyses (Figure 10) demonstrated a general pattern in which larger C pools can lead to higher uncertainty for the corresponding C fluxes (e.g., wood C and autotrophic respiration, leaf C and gross primary productivity, GPP).

In addition to those patterns, some special cases were also identified. For example, the assimilation that excluded SMAP had a lower total C uncertainty and the total C residual error than the full assimilation, likely because of the relatively short residence time and, thus, the short memory of soil water in the modeling. This could be improved by assimilating soil moisture observations more frequently and directly assimilating the water fluxes. This pattern may also be associated with the coarse resolution and shallow soil depth (top 5cm) from the SMAP product. This may be alleviated, at least in part, by downscaling SMAP to a higher spatial resolution by using surface temperature and vegetation indices (Fang et al., 2018) through ML-based algorithms (Xu et al., 2021 and Xu et al., 2022). Similarly, as noted earlier (4.1), the comparison between aggregated Landtrendr AGB and NEON ICs illustrated a scale mismatch between field and remotely-sensed data (the finer the remotely-sensed spatial resolution is, the less accurate it will be) because the NEON vegetation inventory samples over the entirety of each NEON site, which is much larger than that of a single Landtrendr pixel (30 m).

### 4.6 Conclusions

In conclusion, this study demonstrated the significant potential of SDA to inform carbon MRV and carbon cycle science more broadly by utilizing models and multiple data constraints to generate fully harmonized carbon budgets across a wide range of ecosystems, thereby reducing both biases and uncertainties in such C cycle estimates. Our VOI analysis then revealed the relative information contribution of different data sources to these well-constrained ecological variables and processes, quantifying their direct constraints on the variables measured, the indirect constraints on other variables at the same sites, and the borrowing of strength across different spatial locations. Soil carbon was found to be both the most valuable data set for reducing existing uncertainties but also the largest remaining source of uncertainty, highlighting the need for better soil C monitoring systems, better-derived data products that integrate these observations, and improved soil biogeochemical models. These analyses also have the potential to be further used by other agencies or networks (such as NASA, NEON, Ameriflux, etc.), providing the ability to more precisely refine existing observation systems, and to define new systems, by providing the information needed to optimize when, where, and which data types should be collected. Moreover, this study also lays the foundation for valuable opportunities to combine bottom-up land SDA systems with top-down atmospheric studies and bridge the ground-based modeling to the remote sensing platforms at larger and finer spatial scales, which will eventually achieve the complete picture of the general earth surface modeling systems, providing a systematic, accurate, and efficient platform for understanding the general environmental problems (e.g., global climate change).

## Notes

https://github.com/pecanProject/pecan/

### Competing Interest Statement

The authors have declared no competing interest.

### Funding

- NASA CMS: CMS 80NSSC21K0965

