## Supplement text for "Harmonizing terrestrial carbon cycle observations over CONUS NEON sites: Assessing the information contributions of multiple data constraints"

**Supplement table and figure.**

*Table S1. The value of uncertainty increment (in proportion) for carbon/water pools/fluxes of different data assimilation results compared to the results of free run or full run.*

| | Wood C | Leaf C | Soil Water | Soil C | AutoResp | HeteroResp | GPP | Transp |
|---|---|---|---|---|---|---|---|---|
| -AGB | 0.361 | 0.000 | 0.009 | 0.037 | -0.073 | -0.022 | 0.191 | 0.024 |
| -LAI | -0.062 | 0.867 | 0.161 | 0.079 | -0.169 | -0.075 | 0.297 | 0.015 |
| -SMP | 0.069 | 0.019 | 0.732 | 0.005 | 0.121 | -0.040 | 0.091 | -0.001 |
| -SoilC | -0.013 | 0.002 | -0.020 | 0.918 | 0.132 | 0.721 | 0.091 | 0.034 |
| +AGB | -0.390 | 0.002 | -0.017 | 0.123 | 0.009 | 0.072 | -0.126 | 0.010 |
| +LAI | 0.231 | -0.883 | -0.206 | 0.055 | -0.174 | 0.022 | -0.235 | 0.001 |
| +SMP | -0.081 | -0.133 | -0.750 | 0.096 | -0.182 | 0.001 | 0.190 | 0.010 |
| +SoilC | -0.102 | 0.001 | 0.002 | -0.905 | -0.120 | -0.705 | 0.203 | 0.028 |

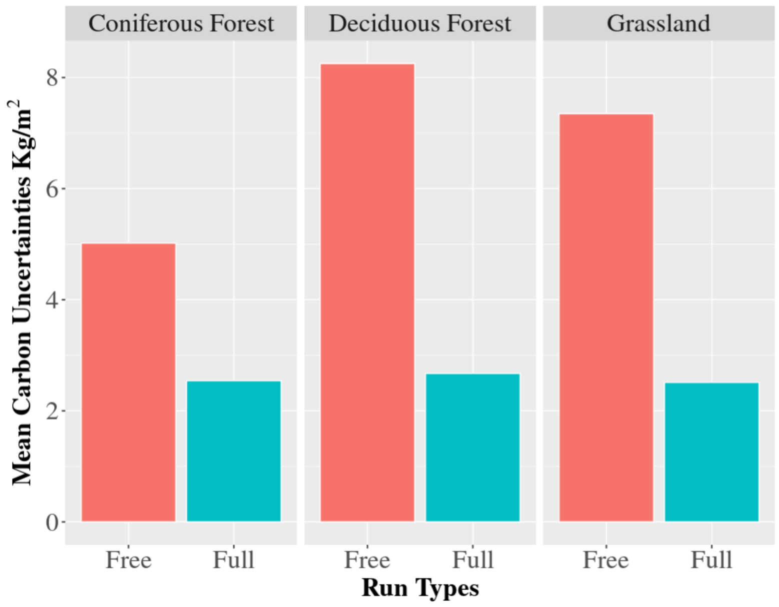


*(A)*

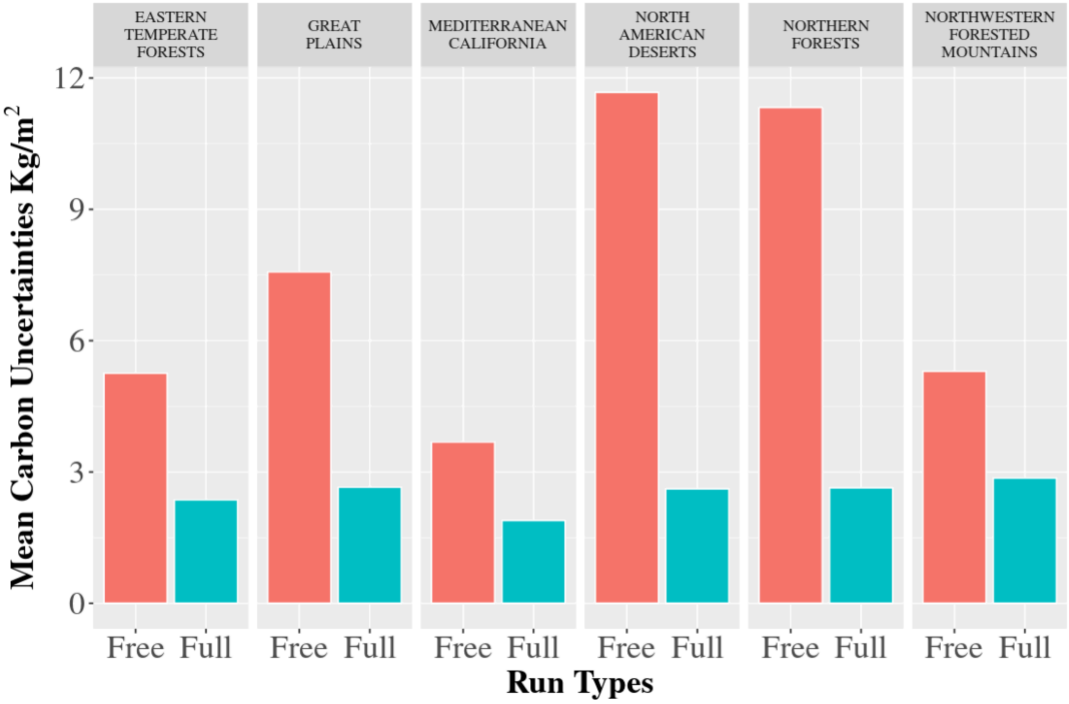


(B)

*Figure S1. The averaged (A) by-PFT and the (B) by-ecoregion carbon uncertainty before and after the SDA. Here, we first summarize the total carbon uncertainties for each NEON site and then average them by PFT or ecoregion.*

**Supplement 1. Initial Conditions from NEON**

SIPNET initial conditions (ICs) represent the initial size of the carbon and water pools. We estimated the ICs of woody, leaf, and soil carbon from NEON measurements. Following Figure S2, we used stem diameter observation (DBH) from the vegetation structure datasets, which took the measurements of trees (DBH >= 10 cm) at squared subplots (10 m) within each plot (400 - 1600 meter square, Meier, 2023) (40 - 50 plots in total) in each site, to calculate the tree-level biomass using equation 1, where a and b are empirical allometric parameters that vary across different tree species (Jenkins et al., 2003), p and t represent the index of plot and tree, while P and T represent the total number of plots in a site and trees within each plot, respectively, and DBH is the observed stem diameter (NEON DP1.10098.001).

$$Biomass_{t,p} = e^{a+b*ln(DBH_{t,p})}$$

$$Tree\ Biomass_{p} = \frac{\sum_{1}^{T} Biomass_{t,p}}{Area_{p}} \quad (1)$$

Tree leaf biomass was calculated using an allometric relationship predicting the proportion of plant biomass that is leaf (equation 2), where the $b_0$ and $b_1$, are empirical allometric parameters in the biomass ratio equation (Jenkins et al., 2003).

$$Ratio_{t,p} = \begin{cases} e^{b_0 + b_0/DBH_{t,p}}, DBH_{t,p} > 2.5m \\ e^{b_0 + b_0/2.5}, DBH_{t,p} \leq 2.5m \end{cases} \quad (2)$$

Plot-level leaf biomass is then calculated by summing the tree leaf biomass over all the trees in the plot and adding the herbaceous dry mass observations (equation 3), where $Herb\ Biomass$ is the herbaceous dry mass observation (NEON DP1.10023.001).

$$Leaf\ Biomass_{p} = (\sum_{t=1}^{T} (Ratio_{t,p} * Biomass_{t,p}) + Herb\ Biomass_{p})/Area_{p} \quad (3)$$

Plot-level woody biomass is just the difference between the total and leaf biomass (equation 4).

$$Woody\ Biomass_{p} = \sum_{t=1}^{T} (1 - Ratio_{t,p}) * Tree\ Biomass_{p}/Area_{p} \quad (4)$$

The NEON soil C datasets are more challenging because SOC estimation requires combining information from the one-time NEON Megapit soil physical and chemical measurements (DP1.00096.001, 0-200cm depth) with the annual plot-level soil physical and chemical measurements from the Periodic datasets (0-30 cm depth) (DP1.10086.001). These two soil chemical observations are combined to ensure that we account for the full soil profile

(0-200 cm) while considering spatial variability and uncertainty across different plots at each site. To do so, we first calculate the soil carbon at different depths (0-30 cm and 0-200 cm) using Equation 5, where the bulk density and carbon concentration are obtained from the NEON Megapit measurements, while the volume is calculated for each depth.

$$SoilC_D = \sum_{depth=0}^{D} BulkDensity_{depth} * Carbon\ Concentration_{depth} * Volume_{depth} \quad (5)$$

After that, we will calculate the fraction of soil carbon between 30 cm and 200 cm depth in equation 6.

$$Fraction_{30cm/200cm} = SoilC_{30cm}/SoilC_{200cm} \quad (6)$$

Then, because the Megapit and Periodic measurements were taken at different depths, we used the averaged bulk density across the entire profile of the Megapit measurements to calculate the SOC at different depths using the Periodic measurements of the organic C percentage (the percentage of organic C in the whole soil matter) for different plots. Finally, the by-plot SOC can be estimated by averaging SOC across depths and dividing by the fraction of SOC between the 30cm and 200cm depths (equation 7).

$$SOC_p = \frac{mean(\sum_{depth=0}^{D} mean(BulkDensity_{0-Megapit\ depth})*OrganicPercentage_{p,\ depth})}{Fraction_{30cm/200cm}} \quad (7)$$

After estimating initial conditions for each plot, we then create ensemble-based initial conditions for each site by resampling plots with replacement and calculating the mean pool size across plots.

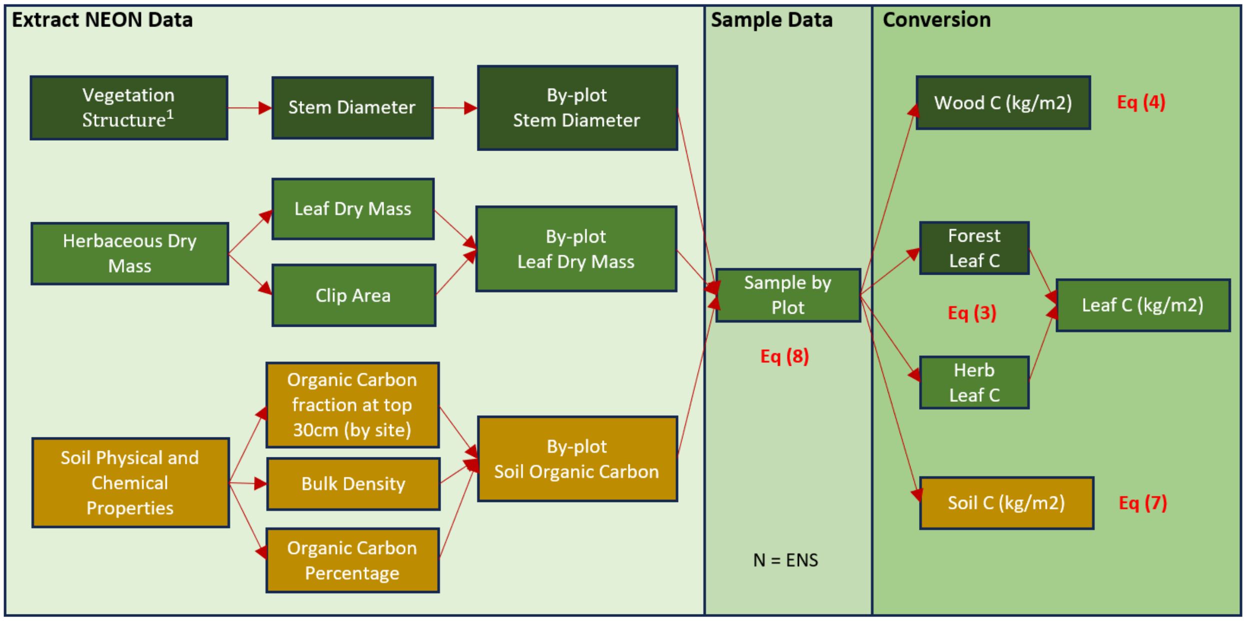

*Figure S2. Workflow for the creation of initial conditions. Here, we summarize the workflow for the leaf, wood, and soil carbon initial pools. NEON data were converted into carbon pool estimates based on equations 1 - 7. After sampling from the measurements by plots or subplots, we finally calculate the carbon density for three carbon pools with proper uncertainties by prescribed ensemble members.*

**Supplement 2. Localization and Block-based Bayesian Sampling**

To improve C cycle estimation and address our research question about shared spatial information, the TGEnF also considers the spatial interactions and correlations among NEON sites, leveraging the fact that nearby sites may share similar ecoclimatic patterns. Within SDA algorithms, the spatial range over which spatial information is shared is controlled by an approach called localization, which sets spatial covariances to zero beyond a specified range. To increase computational efficiency and improve convergence for high-dimensional MCMC sampling, we further developed the localization algorithm by incorporating a block-based data structure design. This approach identifies spatial clusters of interconnected sites and runs distinct clusters separately in the TGEnF.

The recursive clustering algorithm for the localization can be found in Figure S3. Here, we randomly select a site $K$ as the initial site selection. After that, we search the sites $KK$ that are within the spatial scale $S$ to the site $K$ and re-initialize the site selections as $KK$ and so on until the number of initial site selections becomes zero, then we finish the first block with selected sites $K.vec$ and randomly select the rest of the unselected sites to initialize the following block structures until all NEON sites are selected.

Within the iterative forecast/analysis cycle, once we have the forecast results (Figure 4 step 2: $\mu_f$ and $P_f$ for mean and covariance matrix of the ensemble forecasts) and the corresponding observations (Figure 4 step 3: observed data $Y$ and observation error covariance matrix $R$), we split them into small blocks using the clusters calculated during the localization step. Then, because clusters are independent of each other, the MCMC sampling of each cluster is run in parallel on different CPUs, significantly increasing computational efficiency. During the analysis step, we first project the model forecasts into the Tobit space, which is a zero-censored version of the multivariate normal (MVN) that accounts for the fact that our carbon and water pools cannot be negative (equation 9). Process error ($Q_{v,s}$ representing the process error of variable $v$ at the site $s$) is then incorporated using a Normal data model (equation 11) and a gamma prior (equation 10) with parameters $shape_{v,s}$ and $rate_{v,s}$. The updated "analysis" of the model pools, $X$, is estimated using a multivariate Normal likelihood (equation 12). Finally, the ensemble initial conditions are updated using ensemble adjustment based on the analysis mean and covariance (https://github.com/pecanProject/pecan/).

$X.mod \sim MVN(\mu_f, P_f)$ (9)

$Q_{v,s} \sim gamma(shape_{v,s}, rate_{v,s})$ (10)

$X{\sim}Norm(\mu = X.mod,\ sd = 1/Q^{0.5})$ (11)

$Likelihood = MVN(Y|X,\ R)$ (12)

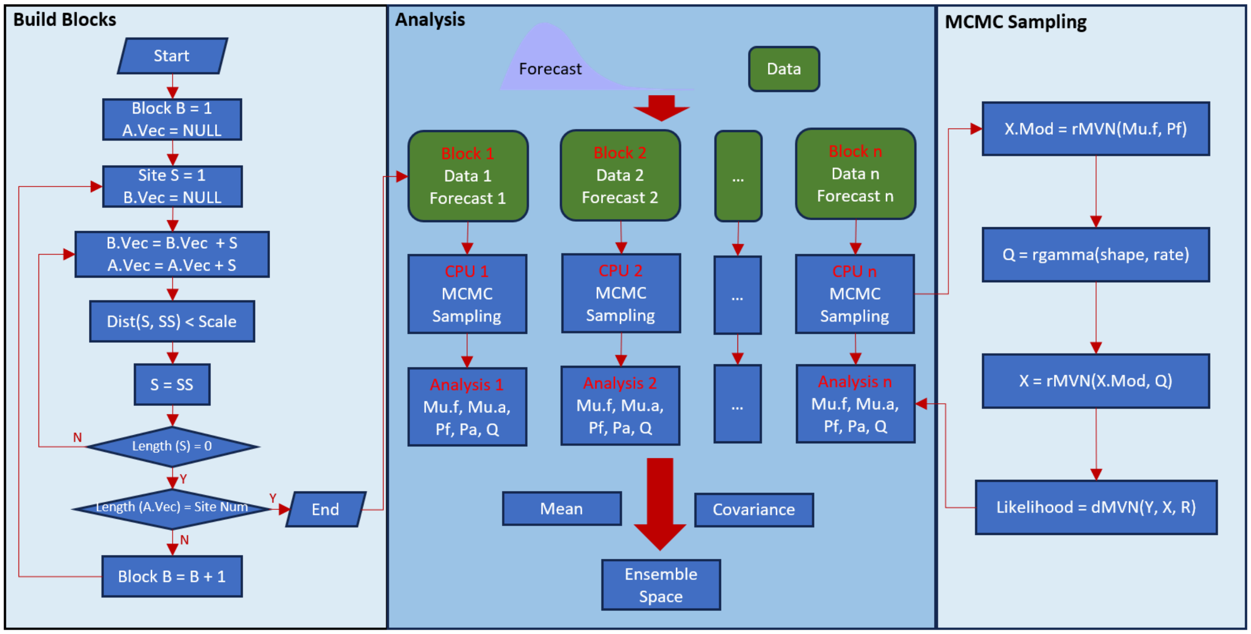


*Figure S3. Workflow of the block-based model-data fusion framework that will be employed in the 4th step in Figure 4, within which we describe the detailed algorithm of how we execute multi-block SDA under the parallel mode. The left panel shows the recursive algorithm for building blocks, executed after the SIPNET model forecast. The middle panel shows the block-based analysis framework, and the right panel shows the algorithm for the Bayesian MCMC sampling.*

### Supplement3. Initial Conditions

In this study, we initiated three carbon pools (leaf, wood, and soil carbon) from the NEON field measurements to start the SIPNET model runs. Discrepancies between remotely sensed and field measurements are well-known in the literature (CITATIONS). Therefore, it is essential to compare the ICs with the remotely sensed observations to ensure we assimilate helpful information into the system.

Figure S4 shows the comparisons between ICs and remotely sensed observations. For the aboveground biomass, the comparisons are generally in good agreement, with an $R^2$ of 0.7 and an overall RMSE of XX (units), except for a few edge cases where Landtrendr showed lower AGB. This mostly occurs when the Landtrendr AGB is near zero for the individual pixel where the NEON tower is located, but the NEON plots capture higher biomass levels at a site scale. Similarly, we found that in a few cases, Landtrendr underestimated biomass when it is at its highest density (~400Mg/ha). However, those edge cases are eliminated when we increase the number of Landsat pixels averaged over (with a maximum of 0.87 $R^2$ between IC and observation at 25 pixels [=150m square]), which suggests that errors are due to the spatial mismatch between remote sensing and field observations (Figure S5). For the soil carbon, the comparison between NEON and SoilGrids shows a similar distribution density while the range of NEON IC is nearly twice as the SoilGrids observations, with the highest NEON SOC values occurring on sites where SoilGrids predicts below average SOC. For the LAI comparisons, while both IC and MODIS LAI share a similar LAI range (from 0 to 7.5), the MODIS LAI tends to be more varied, and the field measurements are concentrated between 0 and 2.5, possibly due to the spatial mismatch between ICs and MODIS LAI observations (Table 1).

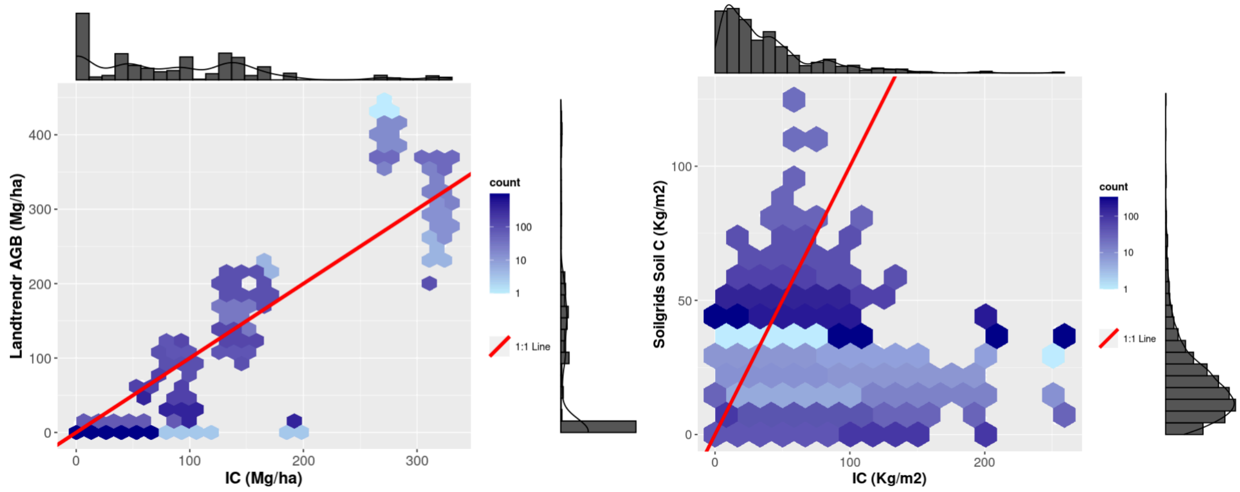

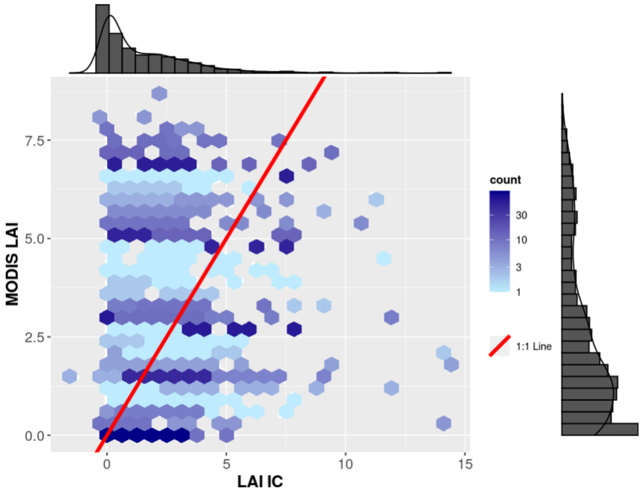


*Figure S4. Comparisons between initial conditions and observations on 2012-07-15. The x-axis represents the ensemble-based initial conditions for 39 CONUS NEON sites, and the y-axis represents the normally sampled values using the mean and variance from observations. The density distributions on the x and y axes represent the marginal density for the initial conditions and observations.*

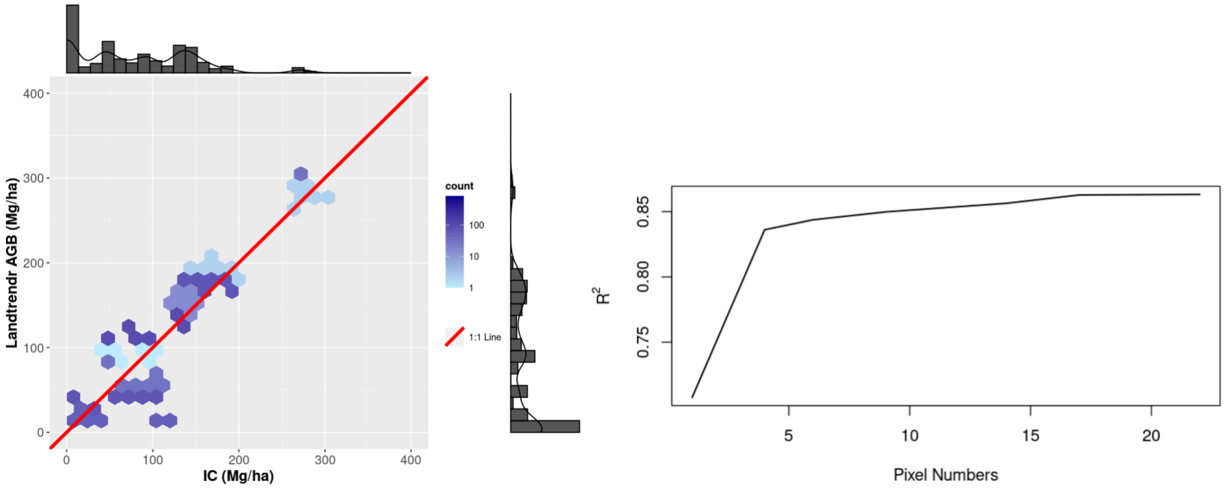


*Figure S5. Comparisons between ICs and Landtrendr observations across the searching radius. The left panel shows the comparisons of woody biomass at a search area of 25 pixels (22550 m²) using the LandTrendr products, which is more accurate compared to the 1-pixel comparisons. The right panel shows the trend of $R^2$ values as the number of pixels averaged increases.*
